# Diverse origins of near-identical antifreeze proteins in unrelated fish lineages provide insights into evolutionary mechanisms of new gene birth and protein sequence convergence

**DOI:** 10.1101/2024.03.12.584730

**Authors:** Nathan Rives, Vinita Lamba, C.-H. Christina Cheng, Xuan Zhuang

## Abstract

Determining the origins of novel genes and the genetic mechanisms underlying the emergence of new functions is challenging yet crucial for understanding evolutionary innovations. The convergently evolved fish antifreeze proteins provide excellent opportunities to investigate evolutionary origins and pathways of new genes. Particularly notable is the near-identical type I antifreeze proteins (AFPI) in four phylogenetically divergent fish taxa. This study tested the hypothesis of protein sequence convergence beyond functional convergence in three unrelated AFPI-bearing fish lineages, revealing different paths by which a similar protein arose from diverse genomic resources. Comprehensive comparative analyses of *de novo* sequenced genome of the winter flounder and grubby sculpin, available high-quality genome of the cunner and 14 other relevant species found that the near-identical AFPI originated from a distinct genetic precursor in each lineage. Each independently evolved a coding region for the novel ice-binding protein while retaining sequence identity in the regulatory regions with their respective ancestor. The deduced evolutionary processes and molecular mechanisms are consistent with the Innovation-Amplification-Divergence (IAD) model applicable to AFPI formation in all three lineages, a new Duplication-Degeneration-Divergence (DDD) model we propose for the sculpin lineage, and a DDD model with gene fission for the cunner lineage. This investigation illustrates the multiple ways by which a novel functional gene with sequence convergence at the protein level could evolve across divergent species, advancing our understanding of the mechanistic intricacies in new gene formation.

## Introduction

Evolutionary adaptation is fundamentally a genetic process, heavily dependent on the emergence of novel genetic components essential for the development of new adaptive traits. New genes, harboring unique functions, stand as a significant wellspring of genetic innovation (Long, et al. 2003; Long, et al. 2013; Santos, et al. 2017), underscoring the need to comprehend the mechanisms governing their origination — a crucial yet comparatively overlooked source of evolutionary innovation. The inaugural paper investigating the origin of new genes, authored by (Long and Langley 1993), marked the beginning of a growing body of research on the evolution of novel genes in the three decades that followed. Initially, gene duplication was proposed as the exclusive method of gene origination (Ohno 1970; Jacob 1977), where the duplicated copy could accumulate mutations and evolve new functions through neo-functionalization or sub-functionalization if the parental gene had multiple functions (Lynch and Conery 2000; Lynch and Force 2000). In recent years, advancements in genomics have increasingly supported alternative routes of novel gene origination, particularly *de novo* birth from genetic material that was previously non-coding for proteins (Tautz 2014; McLysaght and Guerzoni 2015; Schlötterer 2015; McLysaght and Hurst 2016; Schmitz and Bornberg-Bauer 2017; Van Oss and Carvunis 2019). The grand challenge that persists lies in elucidating the molecular mechanisms of new gene origination, deciphering the functions of the proteins they encode, and the fitness the proteins confer (Moyers and Zhang 2015; Guerzoni and McLysaght 2016). In the absence of an ecological context, inquiries related to adaptation are especially challenging to address in model species.

The diverse fish antifreeze proteins are remarkable evolutionary innovations that emerged under strong selective pressures from sea-level glaciation in Polar regions. This makes them ideal systems for exploring the origination and evolution of new genes endowing a crucial adaptive function driven by natural selection. These proteins serve as a quintessential adaptation in marine bony fishes inhabiting frigid polar or subpolar waters, countering the threat of freezing death in icy, sub-zero temperatures. The strong environmental selective pressure of potential freezing death has led to the adaptive evolution in multiple polar and subpolar fish taxa, independently driving the evolution of multiple, structurally distinct types of antifreeze proteins, including antifreeze protein (AFP) types I, II, and III, and antifreeze glycoprotein (AFGP) across various fish lineages. Regardless of structural differences, all AF(G)Ps share the crucial function of protecting the fish from freezing by binding to internalized ice crystals and halting their growths in their blood and body fluids, thus preventing organismal freezing (Devries 1971). Therefore, these proteins definitively represent convergent evolution of a novel function. With well-studied freeze-preventing function, they also exemplify a rare case of a monogenic trait that alone confers a clear life-saving benefit to the organism. The structural variations in AF(G)Ps stem from their distinct genetic ancestry (Cheng and Zhuang 2020). With these unique attributes, fish antifreeze proteins offer excellent avenues into discovering the diversity of possible genetic sources and mechanisms that natural selection has harnessed in evolutionary adaptation.

Over the past two decades, investigations into the evolutionary mechanisms of fish antifreeze protein have significantly advanced the field of evolutionary biology, exemplifying conceptual models and theories related to the emergence of novel genes and functions (Cheng and Zhuang 2020). Our prior work has contributed to the discovery of a clear example of the *de novo* birth of the AFGP gene in northern codfishes (gadids) (Zhuang, et al. 2019). These studies illustrate the formation of essential components for the new gene from noncoding DNA, providing concrete evidence for the "proto-ORF" model (McLysaght and Guerzoni 2015), wherein a nonfunctional ORF (open reading frame) was present before regulatory signals for expression were acquired. We further elucidated the evolutionary process leading to functional innovation of the *de novo* cod AFGP gene and the evolutionary dynamics of the genotype under different strengths of natural selection (Zhuang and Cheng 2021). Other studies on fish AF(G)P evolution have revealed diverse mechanisms underlying new gene origination. These include a rare case of protein sequence convergence between the AFGP gene of the Antarctic notothenioids and the unrelated northern codfishes (Chen, et al. 1997a). The first example of partial *de novo* evolution of the AFGP gene from a functionally unrelated trypsinogen gene in the Antarctic notothenioids (Chen, et al. 1997b; Cheng and Chen 1999). And the neofunctionalization of a cytoplasmic enzyme (sialic acid synthase) into the secreted type III AFP of the zoarcid fishes, supporting the “Escape from Adaptive Conflict (EAC)” model (Deng, et al. 2010). Additionally, the evolution of type II AFP from a pre-existing ancestor C-type lectin (Ewart and Fletcher 1993) provided evidence for transmission to distant species through horizontal gene transfer (Graham and Davies 2021). Lastly, Type I AFP (AFPI) in the starry flounder arose from GIG2 (grass carp reovirus-induced gene 2), with an unrelated function of viral resistance (Graham, et al. 2022). Despite these advancements, among the known fish AF(G)Ps, the evolutionary mechanisms of AFPI are less understood, leaving a notable knowledge gap in the field.

The AFPI is a newly-emerged protein family found in four phylogenetically distant northern marine teleost fish groups –flounders (Duman and DeVries 1976; Graham, et al. 2022), the cunner (Hobbs, et al. 2011), sculpins (Hew, et al. 1980), and snailfishes (Evans and Fletcher 2001). These groups belong to three divergent orders – Pleuronectiformes (flounders), Labriformes (cunner), and Perciformes (sculpins and snailfishes), which divergent from each other approximately 100 million years ago (Hughes, et al. 2018; Near and Thacker 2024). Multiple isoforms of AFPI have been identified in these fishes, expressing in different tissues and varying in protein size and the presence/absence of a signal peptide (Gourlie, et al. 1984; Gong, et al. 1996; Baardsnes, et al. 2001; Evans and Fletcher 2001; Marshall, et al. 2004; Hobbs, et al. 2011). Despite these differences, the protein structures and amino acid sequences of AFPI are remarkably similar across these different fish lineages. Most of them are amphipathic alpha helices comprising 11-amino acid (aa) repeats rich in Alanine (Ala) with evenly spaced Threonine (Thr) residues. Although structurally analogous, suggesting a common ancestry, AFPI genes from different species exhibit distinct intronic sequences and variable utilization of the Ala codon, suggesting a polyphyletic origin instead (Graham, et al. 2013). The emergence of AFPI is thus hypothesized to result from convergent evolution, encompassing both functional and, in some instances, rare protein sequence convergence. This unique characteristic positions AFPI as an exceptional model for investigating diverse evolutionary pathways in the development of a new protein with a novel function, shedding light on the molecular mechanisms underpinning their formation.

In this study, we explored the genomic origins and evolutionary mechanisms of *AFPI* in three of the four AFPI-bearing (AFPI+) taxa. Leveraging the power of cutting-edge long-read sequencing technology, we performed *de novo* whole genome assembly for two species, namely *Pseudopleuronectes americanus* (winter flounder) and *Myoxocephalus aenaeus* (grubby sculpin). Additionally, we included a third AFPI+ lineage, *Tautogolabrus adspersus* (cunner), and 14 other related species from the three taxonomic groups (Table 1). All of these species have chromosome-level genome assemblies, except for the ballan wrasse, which has a scaffold-level assembly available in GenBank. The evolutionary relationships among these species were constructed and visualized with Timetree (supplementary fig.1). We annotated and characterized the complete *AFPI* family and the homologous genomic regions across all 17 genomes. By conducting in-depth comparative analyses, we successfully pinpointed the distinct genetic precursor in each taxon. Furthermore, we deciphered the different evolutionary processes that led to the convergence of protein sequences for this novel gene family in these divergent taxa.

**Table. 1.**
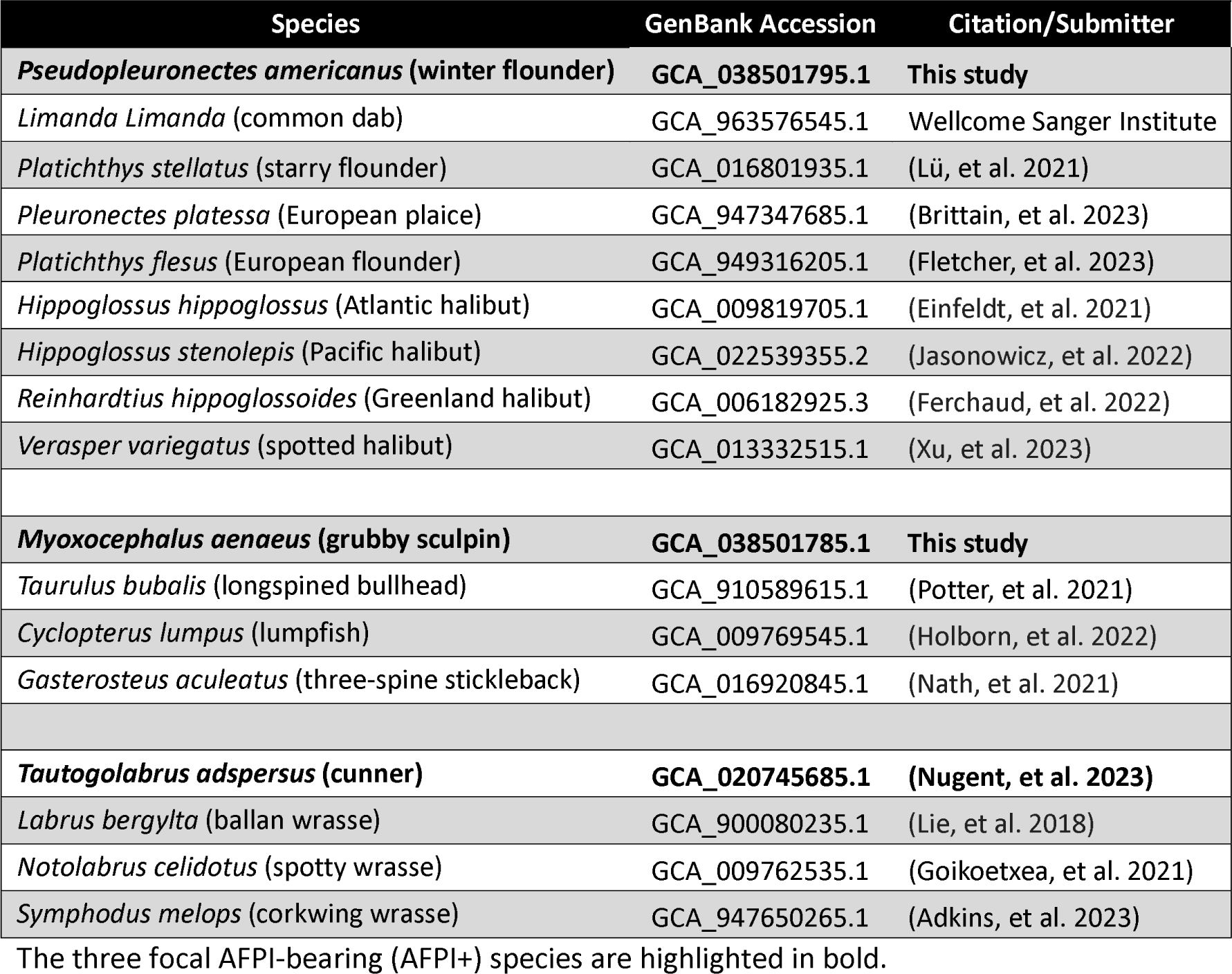
Genome assemblies generated and utilized in this study.

| Species | GenBank Accession | Citation/Submitter |
| --- | --- | --- |
| ***Pseudopleuronectes americanus* (winter flounder)** | **GCA_038501795.1** | **This study** |
| *Limanda limanda* (common dab) | GCA_963576545.1 | Wellcome Sanger Institute |
| *Platichthys stellatus* (starry flounder) | GCA_016801935.1 | (Lü, et al. 2021) |
| *Pleuronectes platessa* (European plaice) | GCA_947347685.1 | (Brittain, et al. 2023) |
| *Platichthys flesus* (European flounder) | GCA_949316205.1 | (Fletcher, et al. 2023) |
| *Hippoglossus hippoglossus* (Atlantic halibut) | GCA_009819705.1 | (Einfeldt, et al. 2021) |
| *Hippoglossus stenolepis* (Pacific halibut) | GCA_022539355.2 | (Jasonowicz, et al. 2022) |
| *Reinhardtius hippoglossoides* (Greenland halibut) | GCA_006182925.3 | (Ferchaud, et al. 2022) |
| *Verasper variegatus* (spotted halibut) | GCA_013332515.1 | (Xu, et al. 2023) |
| ***Myoxocephalus aeneus* (grubby sculpin)** | **GCA_038501785.1** | **This study** |
| *Taurulus bubalis* (longspined bullhead) | GCA_910589615.1 | (Potter, et al. 2021) |
| *Cyclopterus lumpus* (lumpfish) | GCA_009769545.1 | (Holborn, et al. 2022) |
| *Gasterosteus aculeatus* (three-spine stickleback) | GCA_016920845.1 | (Nath, et al. 2021) |
| ***Tautoglabrus adspersus* (cunner)** | **GCA_020745685.1** | **(Nugent, et al. 2023)** |
| *Labrus bergylta* (ballan wrasse) | GCA_900080235.1 | (Lie, et al. 2018) |
| *Notolabrus celidotus* (spotty wrasse) | GCA_009762535.1 | (Goikoetxea, et al. 2021) |
| *Symphodus melops* (corkwing wrasse) | GCA_947650265.1 | (Adkins, et al. 2023) |

The three focal AFPI-bearing (AFPI+) species are highlighted in bold.

## Results

### Genome Assembly and Annotation

We generated a chromosome-level genome assembly of the winter flounder using PacBio CLR (continuous long read) sequencing and Hi-C scaffolding. Our PacBio assembly is highly contiguous, with a contig and scaffold N50 of 1 Mbp and 23 Mbp respectively (Table 2). For winter flounder, 98.73% of the total assembly length was in 24 chromosome-level super-scaffolds, in agreement with the 2n = 48 karyotype previously described for the species (Hoornbeek and Burke 1981). In addition, we also generated a contig-level genome assembly of the grubby sculpin using PacBio HiFi sequencing with a contig N50 value of 4Mbp. Further, Benchmarking Universal Single-Copy Orthologs (BUSCO) analyses revealed a high degree of gene completeness, at 98.1% for the winter flounder and 97.3% for the grubby sculpin (Table 2). Furthermore, we conducted whole-genome annotation for the three focal species (Table 1). In addition to our in-house genome assemblies for the winter flounder and grubby sculpin, we also annotated the genome for the cunner, for which genome data but no annotation was available (Nugent, et al. 2023)number of protein-coding genes predicted for the winter flounder, grubby sculpin and cunner were 24,604, 22,311 and 21,558 respectively.

**Table 2.**
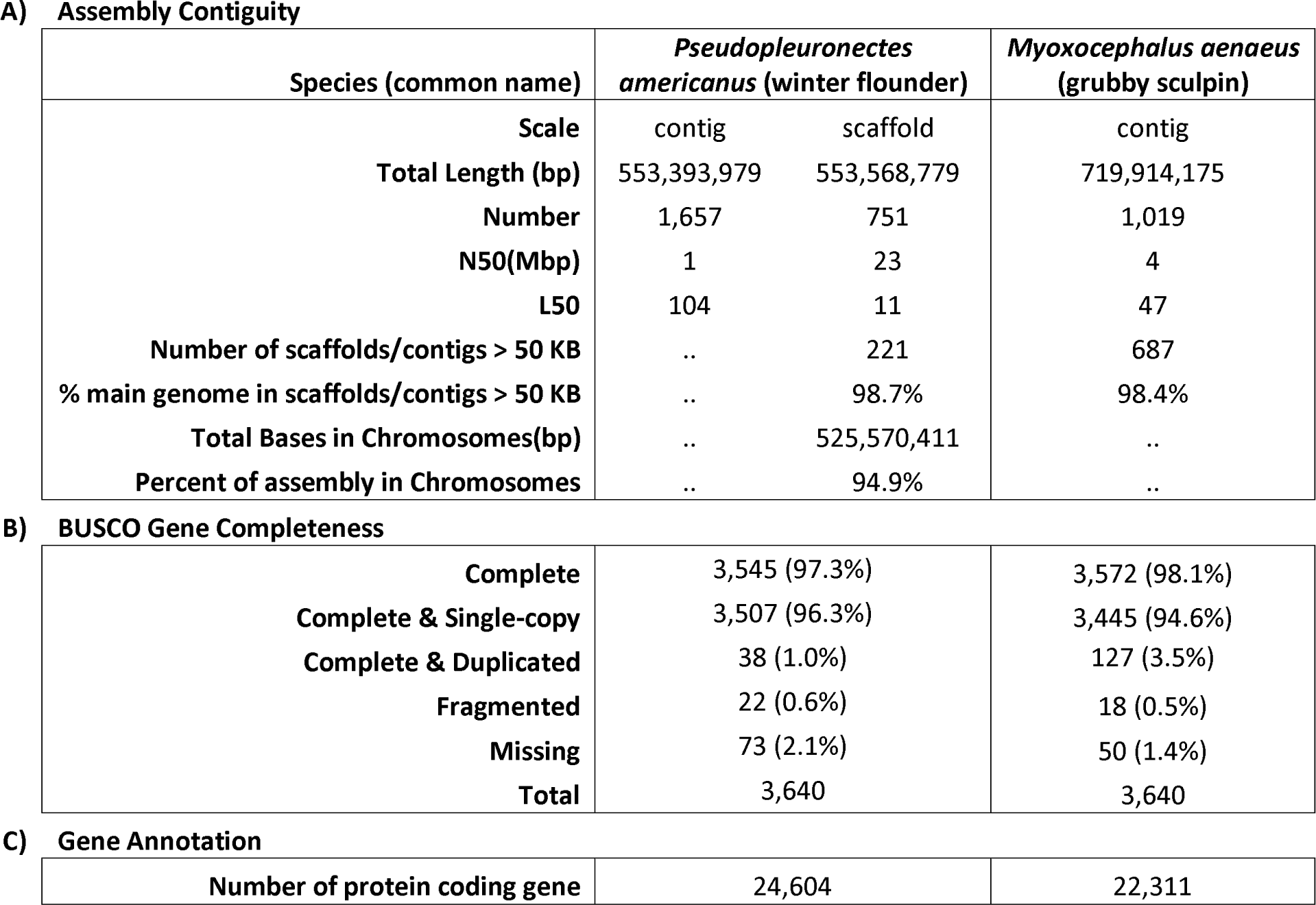
Genome Assembly Assessment.

**A) Assembly Contiguity**
| Species (common name) | *Pseudopleuronectes americanus* (winter flounder) |  | *Myoxocephalus aeneus* (grubby sculpin) |
| --- | --- | --- | --- |
| Scale | contig | scaffold | contig |
| Total Length (bp) | 553,393,979 | 553,568,779 | 719,914,175 |
| Number | 1,657 | 751 | 1,019 |
| N50(Mbp) | 1 | 23 | 4 |
| L50 | 104 | 11 | 47 |
| Number of scaffolds/contigs > 50 KB | .. | 221 | 687 |
| % main genome in scaffolds/contigs > 50 KB | .. | 98.7% | 98.4% |
| Total Bases in Chromosomes(bp) | .. | 525,570,411 | .. |
| Percent of assembly in Chromosomes | .. | 94.9% | .. |

**B) BUSCO Gene Completeness**
|  |  |  |
| --- | --- | --- |
| Complete | 3,545 (97.3%) | 3,572 (98.1%) |
| Complete & Single-copy | 3,507 (96.3%) | 3,445 (94.6%) |
| Complete & Duplicated | 38 (1.0%) | 127 (3.5%) |
| Fragmented | 22 (0.6%) | 18 (0.5%) |
| Missing | 73 (2.1%) | 50 (1.4%) |
| Total | 3,640 | 3,640 |

**C) Gene Annotation**
|  |  |  |
| --- | --- | --- |
| Number of protein coding gene | 24,604 | 22,311 |

### Complete AFPI Genomic Region Characterization in Three Lineages

The completeness of the assembled genomic region and the full set of annotated genes in the genome are pivotal factors in the investigation of new gene origination. We utilized third-generation PacBio long reads, produced using both CLR and CCS (circular consensus sequencing), to capture the contiguous AFPI genomic locus for our analyses. In our assembled genomes, we successfully reconstructed the entire genomic locus that encompasses all members of the *AFPI* family in each species. In addition to defining the complete *AFPI locus*, we characterized the adjacent genomic regions to enable extensive investigation into the evolutionary origins of *AFPI* across three unrelated lineages.

To identify the *AFPI* in each species, we performed BLAST searches using a lineage-specific *AFPI* as the query sequence. All gene members in the *AFPI* family in each species were found in a single locus, fully assembled within a contig of their respective genome assemblies. The locus of winter flounder spans 183 Kbp and contains the entire family of 14 *AFPI* copies. Cunner has 11 *AFPI* in a continuous span of 230 Kbp within one assembled chromosome. Grubby sculpin contains 13 *AFPI* and two additional *AFPI*-like genes within the 347 Kbp *AFPI* locus.

### Distinct Genomic Origins of AFPI Suggested by Absence of Shared Microsynteny and Lack of Nucleotide Sequence Similarity

To investigate the three lineages for potential convergent evolution, we annotated the *AFPI*-containing contigs in the focal species, including all protein-coding genes in their neighboring genomic regions. Our synteny analyses (Fig. 1) revealed that the *AFPI*-containing contigs in the three species do not share microsynteny, *i.e.*, the surrounding genomic and intergenic sequences lack homology among the three species; instead, they display a distinct set of neighboring genes in each species. The absence of shared microsynteny suggests that the *AFPI* in winter flounder, grubby sculpin, and cunner likely have distinct genomic origins. This is attributed to the relatively young age of the gene and the low disruption rate of synteny (Ehrlich, et al. 1997).

**Fig. 1.**
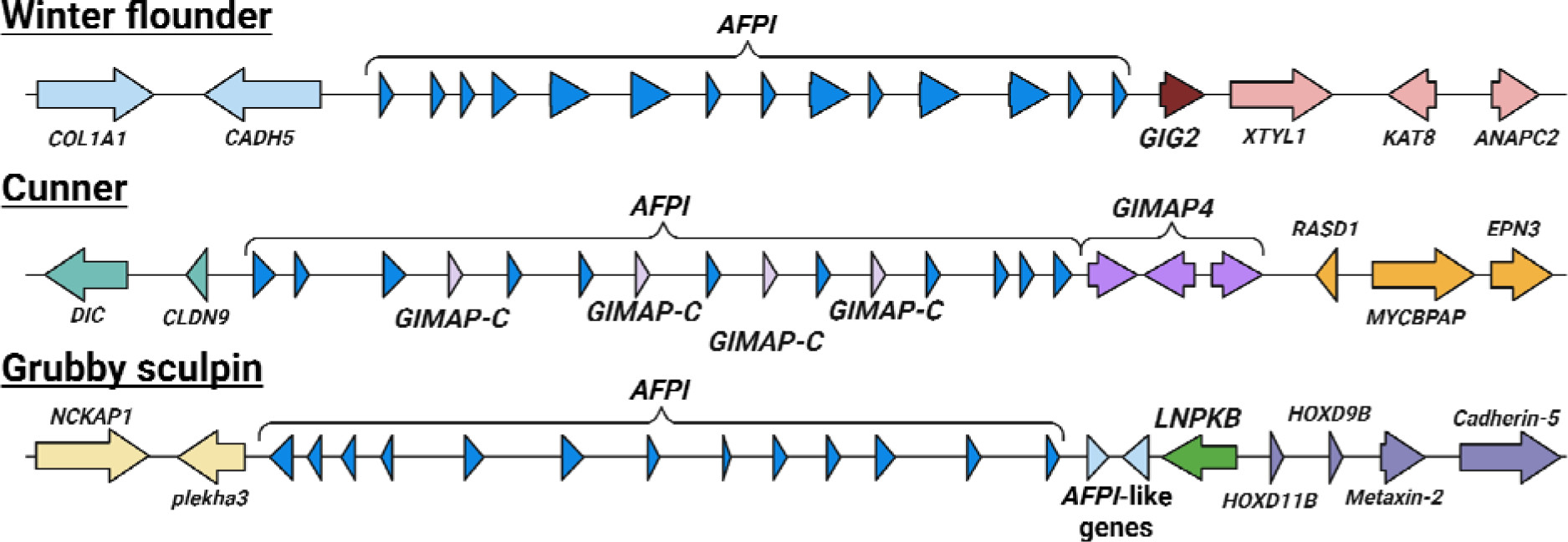
Genomic loci of AFPI and the neighboring genes in the three focal AFPI-bearing species from separate lineages. Arrows and triangles are genes pointing in the sense direction. The AFPI genes are represented in blue triangles, while all unique genes were given a different color.

To further assess the independent evolution of *AFPI* in each lineage, we conducted a multiple sequence alignment and phylogenetic analysis of all *AFPI* across the three species. While their amino acid sequences are very similar (details in the following paragraph), the nucleotide sequences lack homology between the species (supplementary fig. 2). The resulting tree topology of the nucleotide sequences shows that AFPI genes within each species form a separate clade (supplementary fig. 3). The lack of nucleotide sequences homology and the absence of *AFPI* orthologs among the species provides additional evidence for the independent origins of *AFPI*.

### Molecular Convergent Evolution of AFPI Evidenced by Similar Amino Acids and Differential Codon Usage

Despite their distinct genomic origins, the AFPI in these three species exhibits significant sequence similarity. Most share 11-amino acid Ala-rich repeats with evenly spaced Thr residues (supplementary fig. 4), which aligns with the key characteristics of AFPI found in other species (Graham, et al. 2013). Beyond this major common feature related to ice binding, we identify additional commonalities among the AFPI in the three species. In both winter flounder and grubby sculpin, an identical N-terminal motif of ‘MDAPA’ is observed. Likewise, the C-terminal sequence ‘GK*’ is shared between grubby sculpin and cunner (supplementary fig. 4). This additional evidence of amino acid sequence convergence likely serves an important function. The negatively charged aspartic acid (D) in the MDAPA motif at the N-terminus, along with the positively charged arginine (R) or lysine (K) residues at the C-terminus, might be essential for maintaining the stability of AFPI’s helical structural conformation (Harding, et al. 1999).

To further substantiate convergent evolution at the protein sequence level, we conducted an analysis of codon usage preference for *AFPI* within the three lineages (Fig. 2). Expanding beyond the species for which we sequenced genomes, we incorporated additional species with available *AFPI* sequences within each lineage to detect lineage bias rather than species-specific bias. This analysis centered on the Ala residue, as it comprises a significant portion of *AFPI*, ranging from 46.34% to 77.53%. Results suggest that each lineage exhibits a distinct preference for the codon encoding Ala. For example, the flounder lineage shows a preference for the ‘GCC’ codon, the sculpin lineage favors ‘GCG,’ and the only known *AFPI*+ species in the wrasse lineage predominantly utilizes ‘GCT’ (Fig. 2). The discernible codon usage patterns point to distinct genetic origins of *AFPI* in the three lineages, emphasizing the convergent evolution of this novel gene at the amino acid level (Chen, et al. 1997a; Graham, et al. 2013; Athey, et al. 2017).

**Fig. 2.**
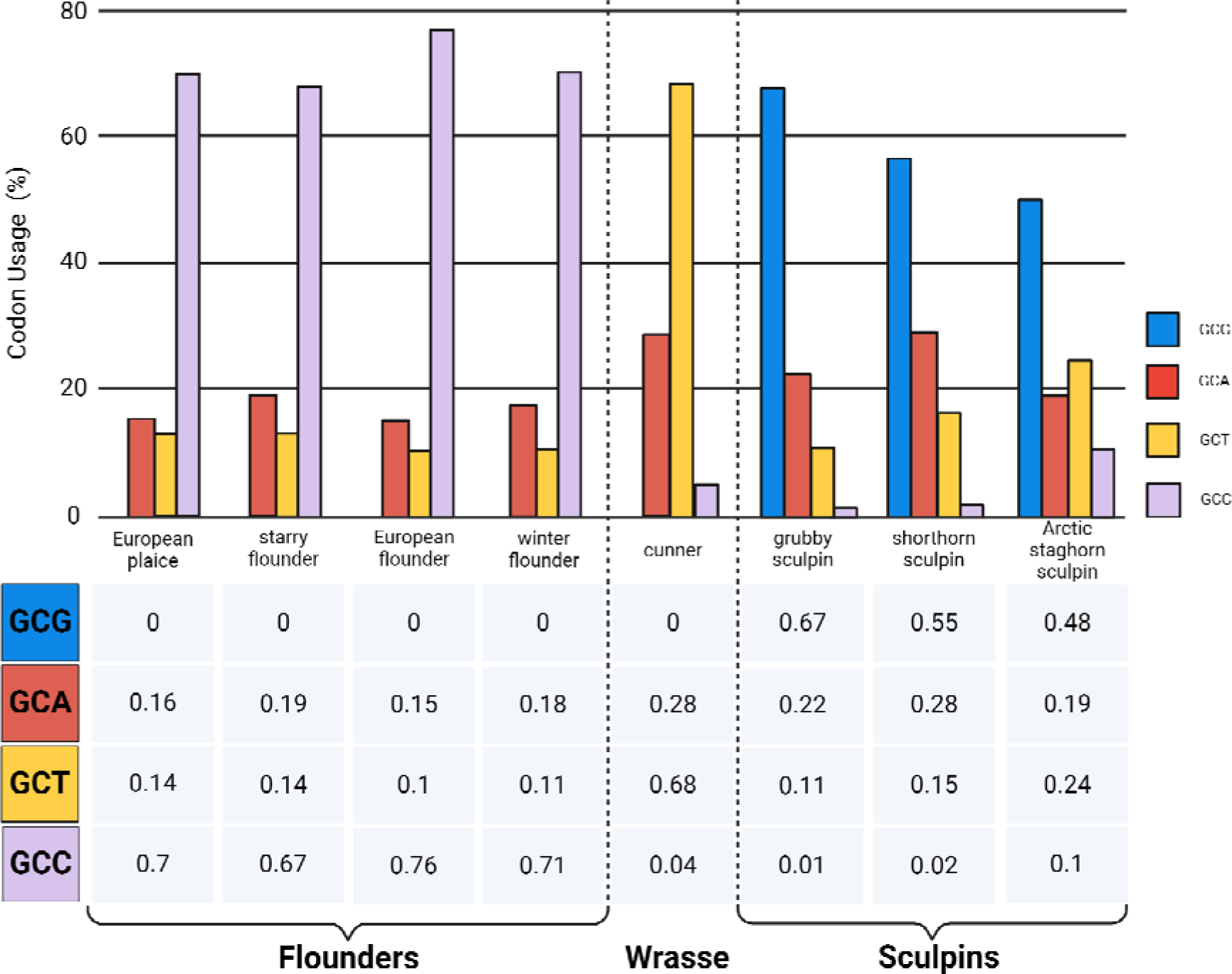
Codon usage for the predominant residue alanine (Ala) across three lineages. In the case of *Pleuronectes platessa* (European plaice), *Platichthys stellatus* (starry flounder), *Platichthys flesus* (European flounder), cunner, grubby sculpin, and winter flounder, AFPI genes were extracted from whole genome assemblies (Table1). For *Myoxocephalus scorpius* (shorthorn sculpin) and *Gymnocanthus tricuspis* (Arctic staghorn sculpin), individual genes were specifically retrieved from GenBank (accession numbers detailed in Material and Methods). Each codon encoding Ala is represented by a unique color.

### *De Novo* Origination of Functional Domains Indicated by Evolutionary Precursors with Unrelated Functions

To identify the extant homolog of the precursor (hereafter referred to as the precursor) of the new gene in each lineage, we thoroughly examined the genomes of both *AFPI*+ and closely related *AFPI*-species. Our approach extended beyond gene sequences, encompassing the entire genome to facilitate homology searching, thereby including any potential non-coding precursor sequences. We found the *AFPI* precursor to be a distinct protein-coding gene in each of the lineages, further corroborating the convergent evolution of this new gene. In the case of the winter flounder, we pinpointed *GIG2* (grass carp reovirus (GCRV)-induced gene 2), which was initially recognized as a novel fish interferon (IFN)-stimulated gene (ISG). In the cunner, we identified *GIMAP4* (GTPase IMAP family member 4-like gene). Additionally, the grubby sculpin precursor was found to function in the endoplasmic reticulum (ER) junction formation, referred to as *LNPKB* (Lunapark-B).

We then performed a fine-scale comparison between each pair of precursor gene and new gene by aligning their corresponding gene components and flanking regions (Fig. 3). For winter flounder, sequence identity spans a significant portion of the *AFPI*, although the coding sequence (cds) lacks homology (Fig. 3A, supplementary fig. 5A). Specifically, 95% sequence identity is observed in the 5’ UTRs and intronic regions between *GIG2* and *AFPI*, providing compelling evidence that *AFPI* originated from a pre-existing GIG2 gene, consistent with a previous study on starry flounder (Graham, et al. 2022). The only cds region sharing sequence identity is a fragment encoding 10 amino acids, corresponding to the sole region in the entire GIG2 protein containing an alpha-helical structure, resembling the structure of the mature AFPI (supplementary fig. 6). This is likely the original coding unit, from which the repetitive cds of *AFPI* was generated through tandem duplication, giving rise to the extended alpha-helical structure responsible for the novel antifreeze function.

**Fig. 3.**
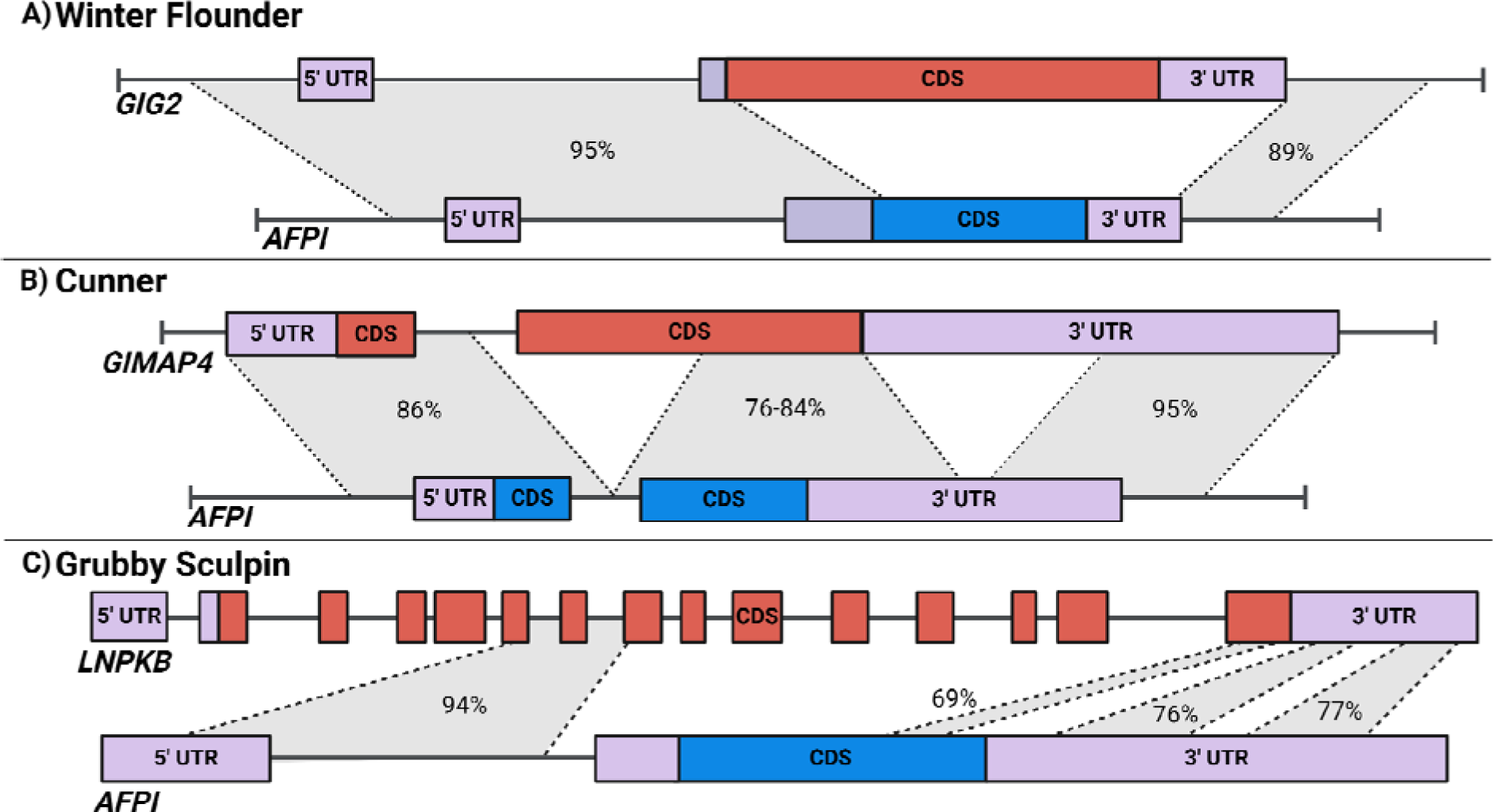
Nucleotide sequence identity analysis using BLAST between ancestral and new genes in (A) winter flounder, (B) cunner, and (C) grubby sculpin. Exons are depicted as boxes, while introns and flanking regions are represented by lines. Untranslated regions (UTRs) are indicated in purple, and the coding sequence (CDS) in the precursor genes and new genes are indicated by red and blue, respectively. Regions exhibiting nucleotide sequence similarities are linked by grey shading, with the corresponding identity percentages.

The precursor gene *GIMAP4* in the cunner shares sequence identity that extends across nearly the entire *AFPI*, with a higher sequence conservation in the UTRs than the cds (Fig. 3B, supplementary fig. 5B). The new gene largely preserves the first exon from the precursor gene, encompassing the 5’UTR and the non-repetitive portion of the cds in the *AFPI*. Mutations introduced a new stop codon for *AFPI*, transforming a segment of the precursor gene’s cds into 3’UTR. Importantly, the duplication of an Ala-rich segment at the C-terminus of GIMAP4 likely played a key role in the development of Ala-rich repeats in AFPI, as evidenced by their common alpha-helical protein structural features (supplementary fig. 6).

The precursor sequence of *AFPI* in grubby sculpin, *LNPKB*, is notably much longer than *AFPI*, spanning over 10 Kbp. By aligning the *AFPI* and *LNPKB* sequences, we discovered a series of homologous regions across the *LNPKB* (Fig. 3C, supplementary fig. 5C). Specifically, the *AFPI* incorporates the 5’ UTR from the 6th exon and the following intron of *LNPKB,* while the 3’ UTR was repurposed from the *LNPKB* 3’ UTR. The cds of *AFPI* shares homology with a repetitive cds found in the last exon of *LNPKB*; while the corresponding amino acid sequences lack similarity, they both feature a similar alpha-helical protein structure (supplementary fig. 6).

The cds of the *AFPI* share reduced similarity or lack similarity with their respective precursor genes in all three lineages. In contrast, their non-coding regions (UTRs, introns, and flanking regions) exhibit high sequence similarity (Fig. 3). The development of the cds with novel antifreeze function of *AFPI* suggests partial *de novo* origination of functional domains in new genes.

### Co-option of Non-Coding Sequences from a Pre-Existing GIG2 Gene formed the Framework for a New AFPI Gene in the Flounder Lineage (family Pleuronectidae)

To reveal the evolutionary process of the new *AFPI* family locus in righteye flounder lineage, we isolated and compared both the *AFPI* and *GIG2* genomic loci, along with their respective homologous genomic regions, from all publicly available high-quality genomes in the Pleuronectidae family (Fig. 4). Among the nine species we examined, we found five are *AFPI*+ and four are *AFPI*-. Each group forms a distinct clade. Notably, winter flounder stands out as having two *GIG2* loci, each located on a different chromosome. The first *GIG2* locus is adjacent to the *AFPI* locus, while the second *GIG2* locus is flanked by the *MTX2* and *BEAN1*. In contrast, *AFPI*- species and the basal *AFPI*+ species *L. limanda* (common dab) only possess the first locus, while the other *AFPI*+ species have only the second locus.

**Fig. 4.**
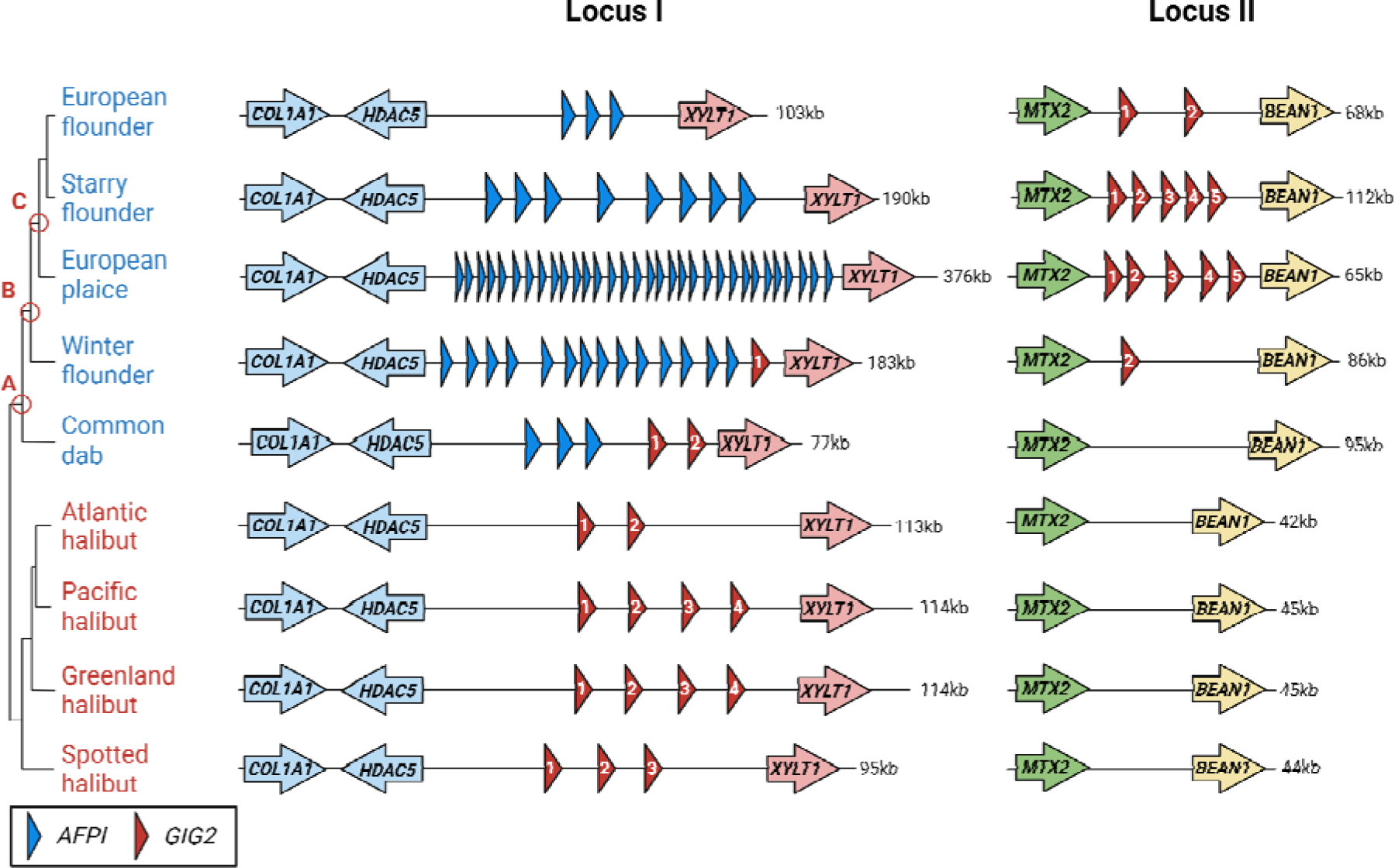
Comparison of *AFPI* and precursor gene locus in *AFPI+* (blue) and *AFPI-* (red) species within the flounder lineage. The circled nodes (A, B, and C) represent the three evolutionary events that shaped the genomic landscape around *AFPI* and *GIG2* locus. Node A displays the origination point for *AFPI* in right-eye flounders. Node B shows the formation of a secondary *GIG2* locus. Node C depicts the loss of *GIG2* in the original locus. The species phylogeny is established from (Vinnikov, et al. 2018). Numeric identifiers within the shapes of *GIG2* are employed to denote the gene names in Supplementary fig. 7.

Based on these patterns, we can deduce the following stepwise evolutionary process (Fig. 4): *AFPI* originated at node A through duplication from a *GIG2* in locus I. Subsequent duplication and/or translocation of a *GIG2* at node B gave rise to a second *GIG2* locus (locus II) located between *MTX2* and *BEAN1*. At node C, there was a deletion of *GIG2* in locus I, accompanied by further duplication of *GIG2* in locus II. Further duplications of the new gene *AFPI* occurred in each species presumably in response to the intensity of freezing selective forces. By incorporating the winter flounder and common dab to represent intermediate forms, we elucidate the step-by-step evolution of the new gene’s origin along the flounder phylogeny. This is further corroborated by the phylogenetic analysis of all *GIG2s* across both loci in the eight species, revealing the *GIG2s* in locus II constitute a distinct clade, with the *GIG2s* in winter flounder and common dab in locus I forming sister clades (supplementary fig. 7). Our inference of this evolutionary process also explains the observed phenomenon in a recent study on starry flounder, where the *AFPI* locus replaces the original *GIG2* locus (Graham, et al. 2022).

### Extension of C-terminal Domain of GIMAP Leads to New AFPI Gene Formation in the Wrasse Lineage (family Labridae)

The *AFPI* locus in the cunner encompasses a series of genes that belong to the GTPases of the Immunity-Associated Proteins (GIMAP) family. Given that cunner is the only known *AFPI*+ species within the wrasse lineage, we isolated the homologous *GIMAP* locus from three related *AFPI*- wrasse species for comparative analysis (Fig. 5). In these three *AFPI*- species, we identified three types of GIMAP genes, namely *GIMAP4, GIMAP7*, and *GIMAP9*. Additionally, we found a different type of *GIMAP*, referred to as cunner-specific *GIMAP* (*GIMAP-C*), which is unique to cunner and has no orthologs in other species (supplementary fig. 8). These *GIMAP-C* are interspersed among the *AFPI* and exhibit different sequences compared to other GIMAP genes.

**Fig. 5.**
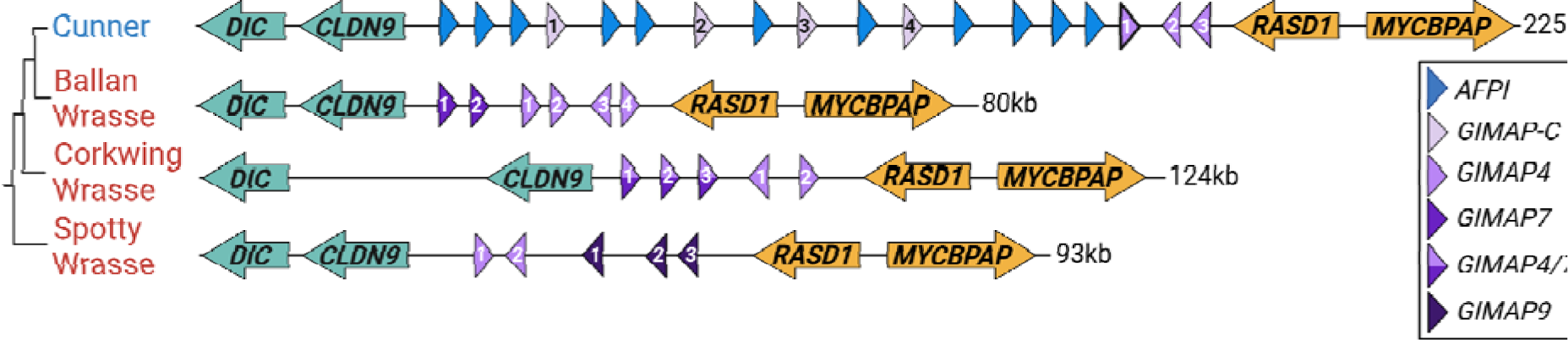
Comparison of *AFPI*/precursor gene locus in *AFPI+* (blue) and three related *AFPI-* (red) species. The phylogenetic relationship between AFPI+ cunner and three related AFPI- species, *Notolabrus celidotus* (spotty wrasse), *Labrus bergylta* (ballan wrasse), and *Symphodus melops* (corkwing wrasse), is based on (Rabosky, et al. 2018). The GIMAP gene annotations were derived from the genome annotations of both Ballan wrasse and Spotty wrasse, with the corresponding accession numbers provided in Table 1. The two-toned shapes represent undefined GIMAP genes that share high sequence identity with both *GIMAP4* and *GIMAP7*. Numeric identifiers within the shapes of *GIMAP* are employed to denote the gene names in Supplementary fig. 8, bolded *GIMAP4* in cunner shares the highest sequence identity with *AFPI*.

Among these different types of GIMAP genes, the first cunner *GIMAP4* (the bolded *GIMAP4-1* in Fig. 5) shares the highest sequence identity with *AFPI*. The GIMAP4 consists of an avrRpt2-induced gene 1 (AIG1) domain followed by a C-terminal region with a stretch of Ala-rich segment and with a Thr residue (supplementary fig. 6). The Ala-rich C-terminus sequence serves as an evolutionary resource for the Ala-rich repetitive of AFPI. While the function of the Ala-rich C-terminus of cunner GIMAP4 remains unclear, the mammalian counterpart of cunner GIMAP4 features an IleGln-rich (IQ-rich) region (Limoges, et al. 2021). Importantly, a 1-nt frameshift mutation can easily alter between Q (Glutamine, codon CAG) and A (Alanine, codon GCA), potentially converting between calmodulin binding function and ice-binding function.

We postulate that an ancestral *GIMAP4* duplicated, giving rise to two daughter genes, one of which evolved into the novel *AFPI*, while the other became *GIMAP-C* (Fig. 6). This hypothesis is supported by the notable sequence identity observed between the precursor and daughter genes. Furthermore, the constructed phylogeny of all available *GIMAP*s across the four wrasse species (supplementary fig. 8) shows that the four *GIMAP-Cs* evolved relatively recently, indicated by their remarkably short branch lengths. They are derived from a *GIMAP4*, forming a sister clade. Subsequent duplications of the newly formed sister gene pair, *AFPI,* and *GIMAP-C*, along with additional *AFPI* duplications, resulted in the observed pattern of *GIMAP-C* arraying with the *AFPI* copies within the locus in the cunner (Fig. 6).

**Fig. 6.**
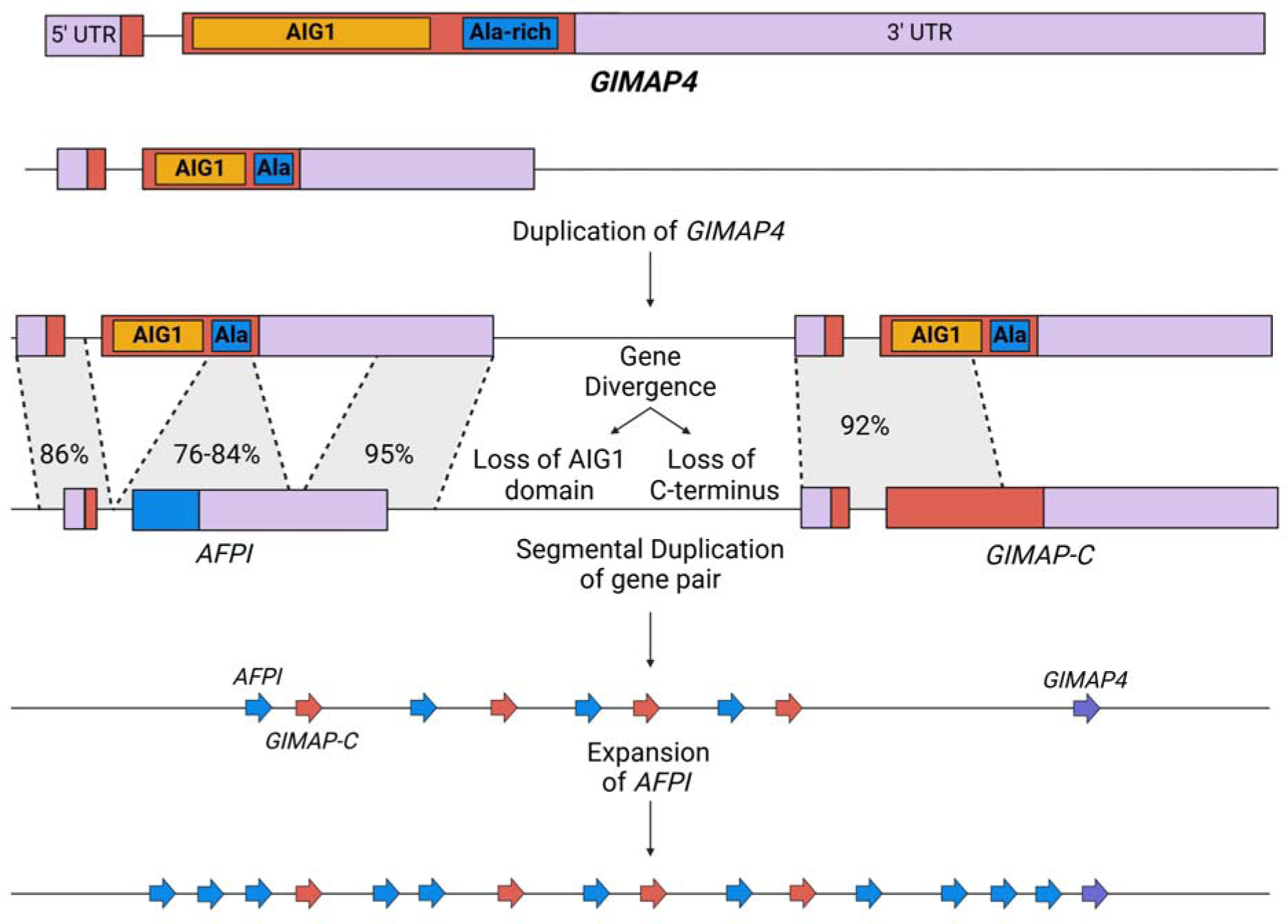
Evolutionary processes of *AFPI* in the cunner. The untranslated regions (UTRs) are depicted in violet, the coding sequence (cds) is highlighted in red, with the AIG1 cds colored in orange, and the Ala-rich cds in blue. *GIMAP4* underwent duplication, and the resulting duplicated genes underwent divergence. The loss of the AIG1 cds led to the emergence of *AFPI*, while the loss of the Ala-rich cds resulted in *GIMAP-C*. These two genes underwent tandem duplication across the locus, with *AFPI* continuing to proliferate.

### Duplication and Degradation in LNPKB Gene Provides Resource for New AFPI Gene Formation in Sculpin Lineage (family Cottidae)

In the case of grubby sculpin, the *AFPI* locus is found in close proximity to the locus of its precursor gene *LNPKB*, which is a single copy gene in both *AFPI*+ and *AFPI*- species (Fig. 7A). In addition, we observed the presence of a fragmented *LNPKB* in the grubby sculpin and its closely related species, *Taurulus bubalis* (long-spined bullhead). In long-spined bullhead, the fragmented *LNPKB* comprises multiple exons, including exon 15 (E15). In contrast, the fragmented *LNPKB* in the grubby sculpin contains most of the parts found in the long-spined bullhead, but lacks E15, while the homologous sequence of E15 is instead found in the new gene *AFPI* (more details in discussion section). The strong sequence conservation observed between the cds part in *LNPKB* E15 and *AFPI* Ala-rich cds suggests a likely origin of the Ala-rich cds within *AFPI* from the *LNPKB* E15. This hypothesis is also supported by the shared protein structural similarity between *LNPKB* E15 and *AFPI*; both exhibit an alpha-helical structure composed of highly repetitive sequences (supplementary fig. 6).

**Fig. 7.**
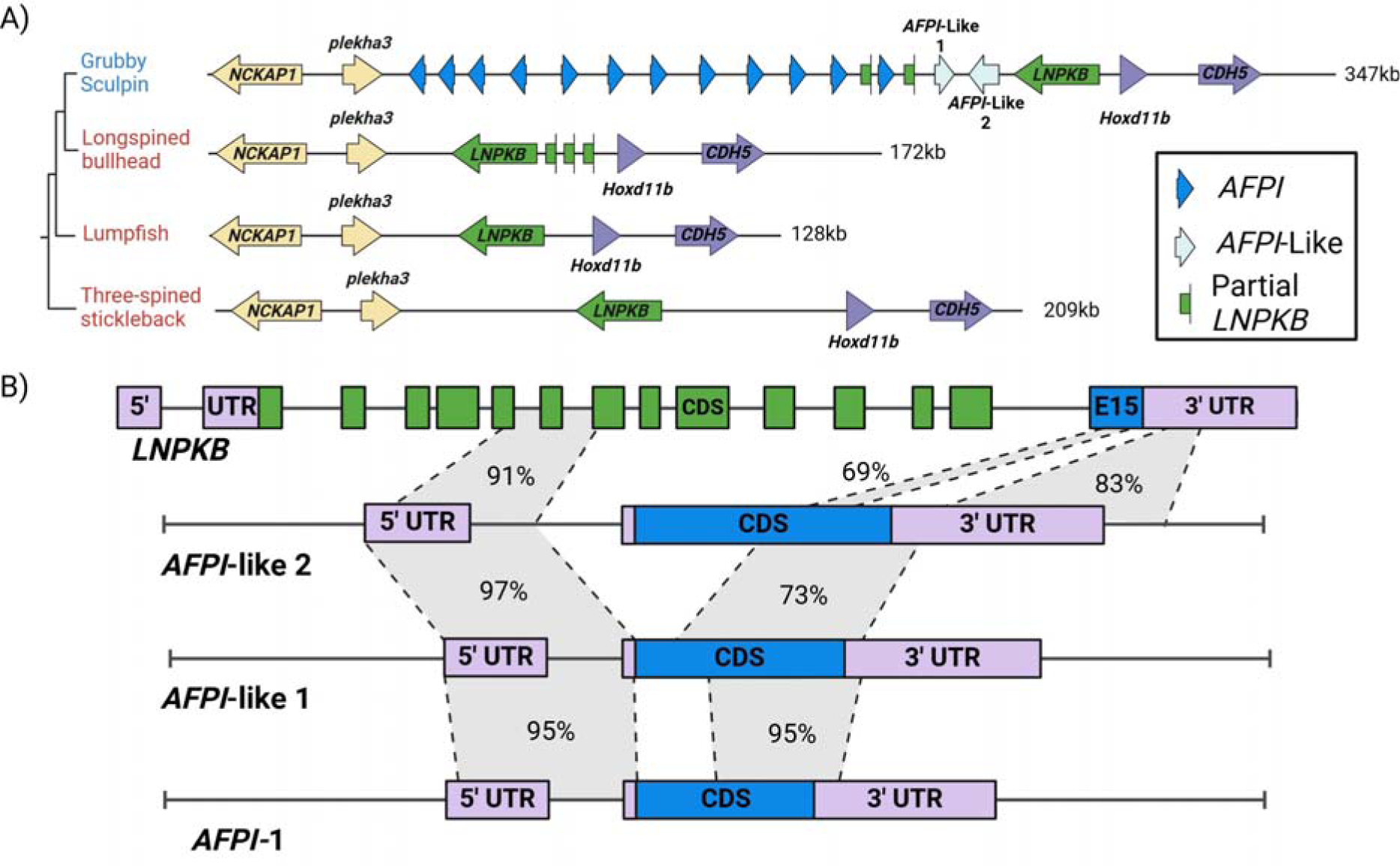
A) Comparison of *AFPI*/precursor gene locus in AFPI+ (blue) and three related *AFPI*- (red) species. The phylogenetic relationship of the *AFPI+* grubby sculpin and *AFPI-* lacking *Taurulus bubalis* (long-spined bullhead)*, Cyclopterus lumpus* (lumpfish)*, Gasterosteus aculeatus* (three-spined stickleback) is based on (Rabosky, et al. 2018). B) Comparison between *AFPI*, *AFPI-like* and precursor gene in grubby sculpin. CDS of *LNPKB* is represented by green and blue boxes, where the CDS in exon 15 (blue) shows sequence identity with the protein CDS in *AFPI*.

To explore the evolutionary trajectory of *AFPI* arising from its precursor gene *LNPKB*, we conducted pairwise alignments of each *AFPI* with *LNPKB* and reconstructed their phylogenetic relationships. This analysis identified two AFPI*-*like genes that share high nucleotide sequence identities with *AFPI* (Fig. 7B, supplementary fig. 5C), but their protein sequences distinguished from AFPIs’ by their N and C terminus, and they feature a repetitive Ala-rich sequence but lack the evenly spaced Thr found throughout AFPI (supplementary fig. 5C). Compared to the *AFPI*, the *AFPI*-like 2 display an extended region (3’UTR) with higher sequence identity to *LNPKB* (Fig. 7B). The AFPI-like 2 contains a repetitive Ala region but lacks the critical Thr, and the AFPI-like 1 contains a couple of interspersed Thr among the Ala-rich repeats, but not evenly spaced as characterized in AFPI. This suggests intermediate evolutionary forms between LNPKB and AFPI, further supported by the phylogenetic analysis (supplementary fig. 9). It appears that these genes represent transitional phases in the evolutionary progression towards refining the 11-aa Ala-rich repeats with evenly spaced Thr to fulfill the ice-binding function observed in AFPI. In contrast to the *AFPI* in the flounder and wrasses lineages, where *AFPI* share the same orientation, the *AFPI* in the sculpin species are oriented differently. The varying orientations of *AFPI*, *AFPI*-like genes, *LNPKB*, and fragmented *LNPKB* in this locus suggest that multiple rounds of inversions and duplications have taken place.

## Discussion

The study of new gene origination is an emerging field in molecular evolution, pivotal for understanding evolutionary mechanisms underlying new traits and adaptive functions. This study investigates the origination of a new gene with known adaptive function in three divergent fish lineages, elucidating the underlying evolutionary mechanisms behind a rare instance of molecular convergence at the protein sequence level. New genes can arise through various mechanisms, including modifying pre-existing genes, sequence rearrangements such as gene fusion or fission, and *de novo* generation of a new Open Reading Frame (ORF) (Long, et al. 2013). The evolutionary pathways of *AFPI* revealed in this study serve as exemplary illustrations of these fundamental mechanisms. Although AFPI in the three lineages evolved from pre-existing genes, their coding sequences do not share homology with these precursor genes, representing a case of partial *de novo* evolution. This unique evolutionary process provides a valuable opportunity to investigate both functional innovations from pre-existing genes and the *de novo* origination of functional domains.

### Evolutionary Models for New Gene Origination

#### Innovation–Amplification–Divergence (IAD) model

While gene duplication is a frequent event in genome evolution, serving as the primary source of material for new genes (Ohno 1970), the occurrence of beneficial mutations creating novel gene functions is rare compared to deleterious mutations that disrupt gene functions. As a result, deleterious mutations often cause a loss of function in one of the duplicated copies long before rare beneficial mutations could occur to drive functional divergence (Bergthorsson, et al. 2007). In support of this, Bergthorsson et al. proposed the **I**nnovation–**A**mplification–**D**ivergence (IAD) model, suggesting that new gene evolution through duplication and divergence begins when a previously irrelevant side activity becomes essential for fitness (**<u>I</u>**nnovation) due to environmental changes (Hughes 1994; Francino 2005; Näsvall, et al. 2012). The initially inconsequential side function gains importance for fitness, and gene duplication (**<u>A</u>**mplification) enhances fitness by increasing the abundance of this initially weak side activity. Then the redundant duplicated gene allows for the improvement of the secondary function under selection, ultimately leading to new gene formation (**<u>D</u>**ivergence).

Our study suggests the origination of *AFPI* in all three lineages can support the IAD model. Alongside their primary functions (e.g. immune or ER-associated), each *AFPI* progenitor harbors an alpha-helical structural motif, with some containing a stretch of Ala-rich sequence (supplementary fig. 6), which provided raw materials for a potentially novel function – ice-binding activity. The decrease in marine temperatures during the late Cenozoic Era (Zachos, et al. 2008; Tripati and Darby 2018) acted as an environmental shift, resulting in the selective pressure that would drive the initially inconsequential sequence segment into a beneficial “**<u>I</u>**nnovation”. Subsequent gene duplication generated a redundant copy, released from the functional constraint of the precursor gene (**<u>A</u>**mplification). The evolution of the duplicate copy continued with the refinement of the Ala-rich repeat sequence and deletion of the precursors’ original coding sequence (**<u>D</u>**ivergence), resulting in the emergence of the new AFPI gene, characterized by 11-amino acid Ala-rich repeats with evenly spaced Thr residues. The new structure imparts strong abilities to bind to ice crystals that invade these cold-water fishes and prevent system-wide ice nucleation.

#### Duplication-Degeneration-Divergence (DDD) model

We propose a new model, the Duplication-Degeneration-Divergence (DDD) model, to depict the formation of AFPI in the cunner and sculpin lineages. The DDD model builds upon the existing Duplication-Degeneration-Complementation (DDC) model (Force, et al. 1999)(Force, et al. 1999) while emphasizing the principles of neofunctionalization rather than the traditional subfunctionalization inherent in DDC. Additionally, the DDD model draws inspiration from the Duplication–Degeneration–Innovation (DDI) model, in which the gene duplication is followed by the differential loss and simplification of the cis-regulatory elements, leading to increased chances for the evolution of new enhancers (Jiménez-Delgado, et al. 2009). In contrast, the DDD model focuses on directly increasing the functionality through the production of novel protein structures. The duplicated copy of the gene must undergo a fate-determining step to be preserved, or it will likely become pseudogenized (Innan and Kondrashov 2010). The DDD model suggests that pseudogenized genes can provide an ideal starting point for the emergence of novel functions. Degenerated genes undergoing coding sequence deconstruction retain essential regulatory elements, allowing for the development of new open-reading frames that bypass the original coding sequence. This process enables a complete divergence from the ancestral gene to the new gene without losing critical regulatory elements. By emphasizing the creation of novel functions, the DDD model accommodates diverse evolutionary mechanisms within its framework. In the following sections, we illustrate how the DDD model applies to the formation of new AFPI genes through different evolutionary pathways in the cunner and grubby sculpin.

In the cunner, the AFPI precursor GIMAP4 belongs to the extensive GIMAP family, which traces its origins back to a time before plants and animals diverged along separate evolutionary paths (Liu, et al. 2008). The primary function of the GIMAP family is related to immunity and it is conserved across a broad spectrum of vertebrates. Within GIMAP4, two crucial domains are identified: the first domain, AIG1, is conserved across all GIMAP4, while the second domain exhibits variations depending on the organism. Notably, in the ray-finned fish clade Percomorphaceae, an Ala-rich second domain evolved in the C-terminus, contrasting with the IleGln-rich domain found in the mammalian counterpart. This distinctive second domain plays an essential role in the origin of *AFPI* when the precursor *GIMAP4* undergoes duplication, degeneration, and divergence, resulting in the emergence of two novel genes.

The DDD model of cunner *AFPI* new gene evolution is facilitated by the evolutionary mechanism of gene fission (Fig. 8). The ancestral gene *GIMAP4* encompasses the major GTP-binding AIG1domain and a secondary Ala-rich C-terminal domain. Through gene **<u>D</u>**uplication, two daughter genes of *GIMAP4* experience **<u>D</u>**egeneration in distinct portions, leading to a single remaining domain in each daughter gene. One daughter gene retains the AIG1 domain, potentially maintaining the GTP-binding function, while the other daughter gene, with the Ala-rich domain, further **<u>D</u>**iverges from the parental gene, enhancing the ice-binding function, and eventually evolves into the new gene *AFPI*. This process aligns with "fission by duplication" (Leonard and Richards 2012), wherein duplicated copies of the ancestral gene lose different domains through either degeneration or separation of the open reading frame. Gene fission was historically deemed uncommon in eukaryotes due to the requisite occurrence of multiple simultaneous evolutionary events at viable positions within each new gene to maintain their expressional machinery (Stechmann and Cavalier-Smith 2002). The DDD model we propose in this study adeptly articulates how one gene can "split" into two, but each daughter gene still retains essential components or regulatory regions such as the promoter, 5’UTR, and start codon (Fig. 8). Supporting this model, gene fission has been documented in the fruit fly *Drosophila*, where two new genes of the *mkg* gene family emerged from the fission of a multidomain ancestor, resulting in two single-domain genes (Wang, et al. 2004).

**Fig. 8.**
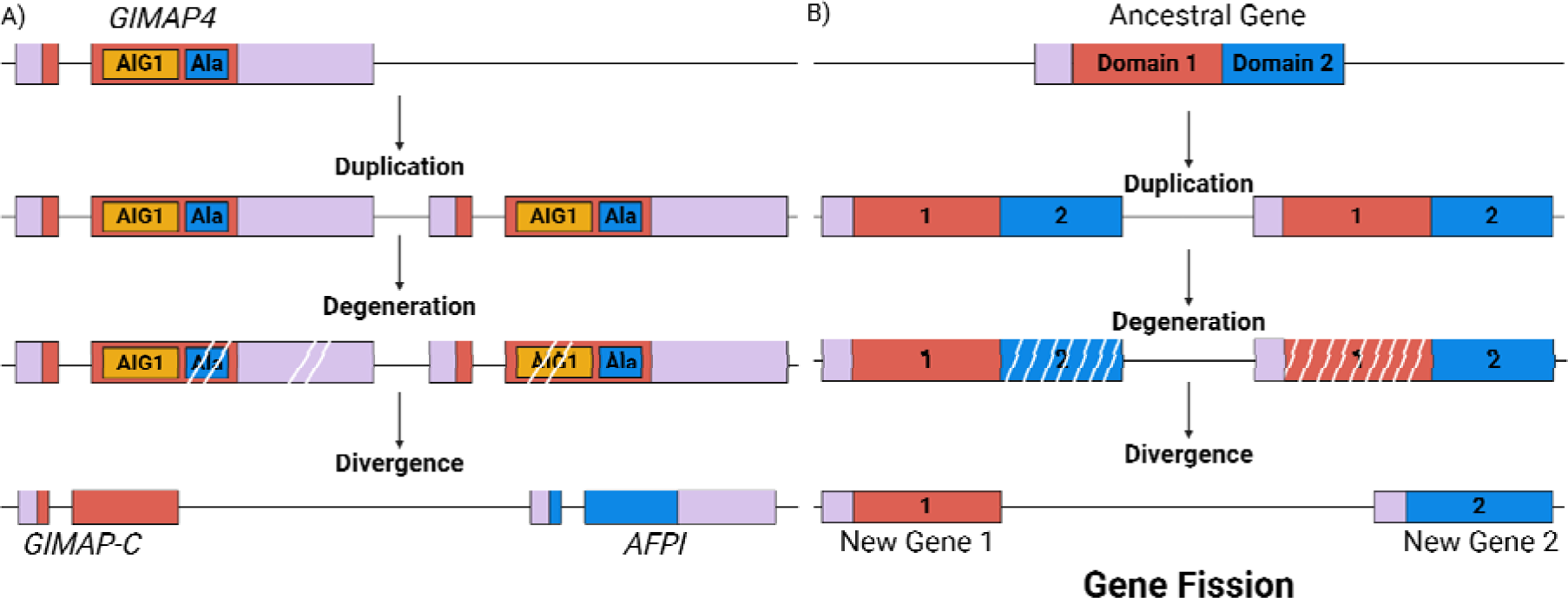
Duplication – Degeneration – Divergence (DDD) model depicting new gene formation in cunner through gene fission. A) The parental gene *GIMAP4* undergoes duplication, degeneration, and divergence to split into the two new genes *GIMAP-C* and *AFPI*. Each new gene inherits a functional domain from the parental gene. B) A simplified illustration of DDD model underlying new gene origination via gene fission. The parental gene initially contains two functional domains. Following gene duplication, degeneration occurs in a different domain in each duplicated gene, and the degraded genes diverged to form two novel, independent genes, both retaining the essential 5’ region.

In the sculpin lineage, the DDD model can be described in a slightly different situation. Near the sculpin *AFPI* family locus, we observed a complete precursor gene *LNPKB* consisting of 15 exons and a fragmented *LNPKB* containing Exons 2-5 and 8-9. Additionally, the two *AFPI*-like genes that represent the intermediate evolutionary stage of *LNPKB* and *AFPI*, exhibit sequence similarity with *LNPKB* Exons 6-7 and 15 (Fig. 7B). Exploring the *LNPKB* locus in the closely related *AFPI*- species long-spined bullhead, we uncovered the presence of one complete and one fragmented *LNPKB* that contained Exons 2-5, 8-9, and 15. Consequently, we deduce the evolutionary trajectory of the new gene *AFPI* in this lineage commenced with the **<u>D</u>**uplication of the *LNPKB*, followed by sequence **<u>D</u>**egeneration of the duplicated gene, and a potential “useful” segment of this degenerated gene subsequently underwent a sequence **<u>D</u>**ivergence, ultimately giving rise to an *AFPI*-like gene (Fig. 9). The *AFPI*-like then underwent further refinement of the new ice-binding function, evolving into an *AFPI*. The *AFPI* evolution in sculpin lineage not only substantiates the DDD model but also lends support to the concept of recycling of genetic material for the creation of novel genes.

**Fig. 9.**
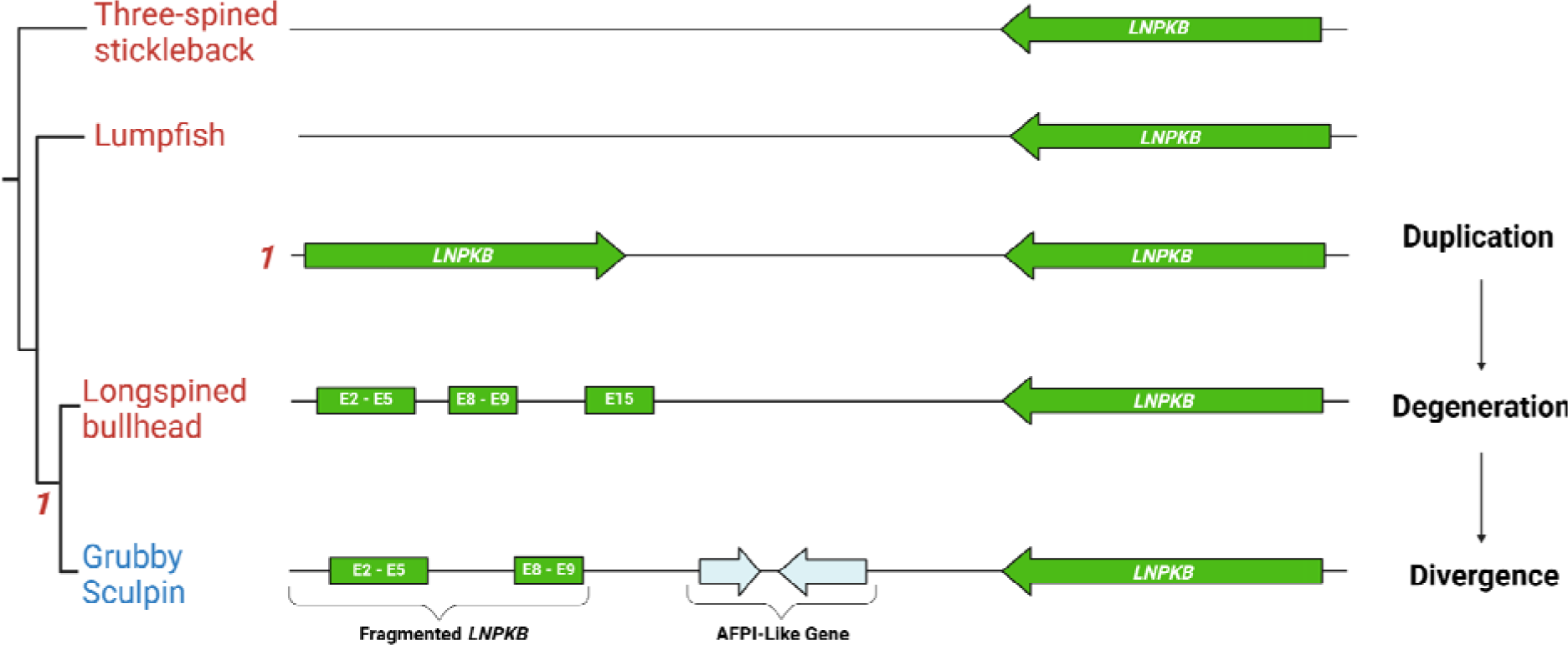
Duplication – Degeneration – Divergence (DDD) model depicting new gene formation in sculpin lineage. The precursor gene *LNPKB* (in green) exhibits a second, degraded copy in *AFPI*+ grubby sculpin and its closely related *AFPI*- species, long-spined bullhead. Fragments of the degraded copy further diverge, giving rise to an *AFPI*-like gene (depicted in blue). A deduced intermediate stage of *LNPKB* duplication is indicated with a 1.

### Evolutionary Drivers Behind New Gene Formation: Cis-Regulatory Elements or Protein Structure Change

Evolution of novel gene function relies heavily on the changes in coding sequences underlying protein structure and the alterations in cis-regulatory elements controlling gene expression. However, the relative contributions of these factors to trait evolution have historically been controversial (Carroll 2005; Hoekstra and Coyne 2007). The perspective that underscores evolutionary changes in anatomy and lifestyle are more frequently rooted in alterations to the mechanisms controlling gene expression than in sequence changes in proteins was initially proposed in 1975 (King and Wilson 1975). This notion has since been substantiated by numerous empirical studies, particularly within the field of evolutionary developmental biology (evo-devo) (Carroll 2005, 2008). Furthermore, many examples of adaptive *cis*-regulatory mutations tend to focus on trait loss rather than gain (Hoekstra and Coyne 2007). Instances include skeletal armor in three-spine sticklebacks (Shapiro, et al. 2004), pigmentation on *Drosophila* wings (Gompel, et al. 2005; Prud’homme, et al. 2006), and dorsal bristle density on *Drosophila* larvae (Sucena and Stern 2000).

In contrast, with regard to trait gain and new gene formation, our study shows a critical role of changes in coding sequences in adaptive evolution via neofunctionalization. The new gene, *AFPI*, in each lineage inherits the non-coding part of the gene framework from the precursor gene, including UTRs/introns, and some flanking sequences. However, the essential ice-binding motif with the adaptive novel function — 11-aa Ala-rich repeats with evenly spaced Thr — is exclusively present in the new gene *AFPI*, not in any precursor genes. This suggests an adaptive strategy for new gene formation, preserving inherited regulatory sequences while developing a novel coding sequence that facilitates attaining a new function. Similar situations were found in type III AFP in Antarctic eelpout (Deng, et al. 2010) and antifreeze glycoprotein (AFGP) in Antarctic notothenioids (Chen, et al. 1997b), where new genes recruited major regulatory sequences (e.g., UTRs) from precursor genes while developing coding sequences with novel functions through substantial sequence alterations or partially *de novo* origination. In fact, all *de novo* genes, arising from ancestral non-coding sequences to form new coding sequences, stand as evidence supporting the perspective of structural adaptation, underscoring the role of coding sequences in fueling evolution.

To further explore the evolutionary dynamics governing structure and regulation of this new gene, we investigated the potential functional or expressional connections with the respective precursor in each lineage. Intriguingly, the precursor genes in two of the three lineages exhibit immune-related functions, specifically flounder *GIG2s* and cunner *GIMAPs*. Genes involved in immune and defense response are generally expected to be more diverse due to the selective pressure imposed by pathogens (Enard, et al. 2016). Therefore, we deduce that the diversity and adaptability inherent in these immune gene families play a crucial role in facilitating the origin of the new gene *AFPI*. For instance, *GIG2*s within the flounder lineage have experienced dynamic evolution, evident in varying copy numbers and distinct genomic loci among closely related species (Fig. 4). In the case of *AFPI*, given the highly repetitive coding sequence, the development of a functional ice-binding peptide through duplication from a short stretch of Ala-rich and Thr-containing sequences is conceivable. However, new genes must acquire a specific transcriptional regulatory system to ensure certain temporal and spatial expression patterns. To some extent, the expression of antifreeze function in response to invasive ice crystals may parallel the induction of immune function in response to invasive pathogens. Nevertheless, this hypothesis awaits future testing using gene expression data. AFPI offers a unique system for studying different paths of new gene formation due to its independent emergence in three divergent lineages. This system also provides an intriguing perspective for future research on how new genes integrate into existing regulatory networks.

## Materials and Methods

### *De Novo* Genome Sequencing and Assembly

Specimens of winter flounder and grubby sculpin were collected from Long Island, New York. High molecular weight DNA was extracted from blood cells embedded in agarose blocks using the Nanobind CBB kit (Circulomics/PacBio). Construction of PacBio SMRT-bell sequencing libraries and long read sequencing on PacBio Sequel II were conducted at the Roy J. Carver Biotechnology Center, University of Illinois, Urbana-Champaign. For winter flounder, PacBio CLR sequencing on one SMRT cell (30 hours of sequence capture) was performed. PacBio HiFi sequencing on one SMRT cell (30 hours of sequence capture) was carried out for the grubby sculpin generating the highly accurate CCS (circular consensus sequencing) reads. The sequencing yielded a total of 14.89 million CLR reads for the winter flounder, totaling 166.9 Gbp, equivalent to approximately 238X genome coverage based on an estimated genome size of 700 Mbp from the Animal Genome Size Database (http://genomesize.com/). For the grubby sculpin, 1.34 million CCS reads totaling 18.7 Gbp were obtained. This is equivalent to about 21.5X genome coverage, using the estimated genome size of 870 Mbp of the closely related shorthorn sculpin (Hardie and Hebert 2003), since the grubby sculpin genome size is unavailable in the database. Additionally, the genome coverage can also be estimated as 26X based on the assembled genome length (Table 2).

To optimize the assembly of winter flounder CLR reads, we compared three genome assembly tools: CANU v2.2 (Koren, et al. 2017), FLYE v2.7 (Kolmogorov, et al. 2019), and WTDBG2 v2.5 (Ruan and Li 2020). These tools were applied to subsets of raw read data covering various read length distributions and depths of coverage following published subsampling strategies (Rayamajhi, et al. 2022). Assembly quality was assessed using BUSCO v3.0.1, utilizing the "actinopterygii" reference gene set (Simão, et al. 2015). The assembly yielding the highest contiguity and gene completeness was obtained by employing FLYE v2.7 with the entire raw read dataset. Grubby sculpin’s CCS HiFi reads were assembled using HIFIASM v0.15 (Cheng, et al. 2021) using default settings.

### Hi-C Genome Scaffolding

For scaffolding the genome assembly, a Hi-C chromosome conformation capture library was prepared for winter flounder using the Proximo Hi-C library kit, and sequenced on the Illumina NovaSeq6000 platform, generating 257.8 million paired-end reads of 150 bp each. The Hi-C-based scaffolding tool SALSA2 program (Ghurye, et al. 2019) was employed for contig scaffolding. Hi-C reads were firstly mapped using the Arima-HiC mapping pipeline (https://github.com/ArimaGenomics/mapping_pipeline). After removing the 3’-side of chimeric mappings, paired BAM files were merged and converted to BED format using SAM tools v1.12.33 (Li, et al. 2009; Li 2011) and BED tools v2.30.0 (Quinlan and Hall 2010). The scaffolding process utilized SALSA2 v2.335 with parameters -e GATC -m yes. To validate and enhance assembly at the chromosomal level, we also applied the YaHS scaffolding tool (Zhou, et al. 2023), which generated more contiguous scaffolds. To address any potential assembly errors, Hi-C contact maps were generated using Juicer v3.0. These maps were meticulously visualized, and manually curated using Juicebox version 1.11.08 (Durand, et al. 2016). These steps involved the removal of residual duplicate contigs and the correction of mis-joins, ultimately leading to a refined chromosome-level genome assembly.

### Genome Annotation

We performed genome annotation using both repeat-masked and un-masked genome sequences. In the first round of annotation, we chose to omit the standard practice of repeat-masking due to the potential masking of *AFPI* repetitive cds, which are crucial for our study. For the second round of annotation, repeat elements were annotated by building a *de novo* repeat library with the RepeatModeler v2.0.3 pipeline (Flynn, et al. 2020), using the BuildDatabase option and the NCBI database as input, and then the repeat masking step was completed using RepeatMasker v4.1.3 (Smith, et al. 2013). The un-masked annotation provided a more comprehensive annotation for the exons of highly repetitive *AFPIs*. Conversely, the masked annotation overlooked repetitive *AFPI* cds, yet managed to predict gene margins with accuracy for other protein-coding genes. We used lineage-specific protein sequences as references: *Hippoglossus hippoglossus* (Atlantic halibut) for flounder lineage, *Cheilinus undulatus* (humphead wrasse; GCA_018320785.1) for cunner, and *Gasterosteus aculeatus* (three-spined stickleback) for grubby sculpin. In flounder lineage, where the *AFPI* precursor *GIG2* family occupies distinct genomic loci outside from *AFPI* locus, besides winter flounder, we also annotated the whole genomes of selected *AFPI*+ and *AFPI*- representatives from each clade, including starry flounder, Greenland halibut, and spotted halibut (Table 1). The ProtHint protein mapping pipeline (Brůna, et al. 2020) was employed to generate required hints from each reference. Subsequently, the BRAKER2 V2.1.6 pipeline (Brůna, et al. 2021) with both GeneMark-ET and AUGUSTUS was used to perform gene prediction. The assembled scaffolds, along with hints generated from the reference protein were utilized to create initial gene structures through the GeneMark-ET tool (Lomsadze, et al. 2014). The initial gene structures were then used to train AUGUSTUS to produce the gene predictions (Stanke and Waack 2003). The final gene prediction resulted from the union of both AUGUSTUS and GeneMark-ET predictions. Additionally, the predicted genes from the BRAKER2 pipeline underwent assessment through BUSCO V5.5.0 (Manni, et al. 2021) to evaluate the completeness of the assembled genome, utilizing the actinopterygii_odb10 database. All GenBank Accession numbers can be found in Table 1.

### Characterization of AFPI Gene Family

We systematically identified *AFPI* across all examined genomes and manually annotated them in *AFPI*+ species. Utilizing available cDNA sequences from GenBank (winter flounder: X07506.1, cunner: JF937681.2, and grubby sculpin (AF305502.1, MH745497.1), we performed homology searches via blastn in BLAST+ tool (Camacho, et al. 2009), excluding the low complexity filter due to the repetitive nature of *AFPI*. Given length variations in the Ala-rich repetitive region among *AFPI*, we performed an additional search by excluding the Ala repeat from the *AFPI* cDNA queries. Subsequently, we annotated the BLAST hits onto the *AFPI*+ contigs using SnapGene software (www.snapgene.com), a nucleotide visualization and annotation tool. The *AFPI* were manually annotated for their intron/exon boundaries and untranslated regions (UTRs). Tandem Repeat Finder (Benson 1999) was used to detect if an Ala repeat motif was present in each gene.

### Codon Usage Bias Analysis

We computed the codon usage for Ala in the coding sequence of *AFPI* within *AFPI*+ representatives (Fig. 2) across each lineage using the Sequence Manipulation Suite: Codon Usage (Stothard 2000). Specifically, within the flounder lineage, we examined all four *AFPI*+ species (winter flounder, European plaice, starry flounder, and European flounder) with available genome data. In the wrasse lineage, the only known *AFPI*+ species, cunner was used in the analysis. In the sculpin lineage, as no other high-quality genome assembly is accessible aside from grubby sculpin that we sequenced, we obtained *AFPI* sequences from GenBank for three additional *AFPI*+ species: shorthorn sculpin (Hew, et al. 1980) (AF305502.1), longhorn sculpin (Low, et al. 2001) (AF306348.1), and Arctic staghorn sculpin (Yamazaki, et al. 2019) (MK550897.1). Due to the strikingly similar codon composition patterns exhibited by all *AFPI* within the same species, one gene per species was selected for comparative analysis. The percentage of Ala-encoding codons was then calculated for each species.

### Characterization of AFPI Family Genomic Locus and Flanking Regions

To delineate the neighboring genes and intergenic sequences around the *AFPI* family, we isolated the scaffolds containing *AFPI* in each *AFPI*+ species from their respective genome assemblies. The BRAKER2 annotations for each sequence were imported into SnapGene Viewer and underwent manual editing. All annotated genes were validated through BLAST searches. In cases where BRAKER2-annotated nucleotide sequences could not be verified using BLASTN, the corresponding amino acid sequences were queried using BLASTP against the non-redundant protein sequence database. The intergenic sequences underwent scrutiny for potential missing annotations using BLAST.

### Characterization of Syntenic Regions in AFPI-lacking Outgroup Species

Upon the annotation of all protein-coding genes in the *AFPI* locus and its flanking regions in *AFPI*+ species, we proceeded to identify syntenic regions in related *AFPI*- species by anchoring onto the two flanking genes of the *AFPI* locus. Specifically, we employed BLAST searches to locate the orthologs of those neighboring genes in *AFPI*- species. Subsequently, the extended syntenic regions in *AFPI*- species were annotated using the same procedure described above for *AFPI*+ species. For Greenland halibut, spotted halibut, and long-spined bullhead, in-house annotation was utilized, while publicly available annotation files were employed for other *AFPI*- species, with manual corrections incorporated. For example, in Atlantic halibut and Pacific halibut, two adjacent GIG2 genes were initially misannotated as a single gene, and we rectified this by reannotating them as separate genes.

### Identification of Ancestral Sequence in AFPI-bearing Species

We searched for the potential ancestral sequence of the new gene *AFPI* in the genome of both *AFPI*+ and *AFPI*- species within each lineage using blastn in BLAST+ tool (Camacho, et al. 2009). Given that the ancestral sequence may not necessarily reside in the *AFPI* locus and might not be a protein-coding gene, a thorough exploration of the entire genome became essential. For the BLAST search, we employed the complete *AFPI* and the non-repetitive segment of the gene as separate queries. Given the highly repetitive nature of the coding region of *AFPI*, we opted to disable the low complexity filter for the BLAST when using the complete gene sequence as a query. Loci containing sequences exhibiting similarity to *AFPI*, but not the members of *AFPI* family, were then isolated to confirm the homology with *AFPI*.

To verify the homologous ancestral sequences, we performed additional pairwise alignments between the potential ancestral sequence and *AFPI* in each lineage. When the precursor gene belongs to a gene family, we annotated all members within that gene family, and isolated the gene sequences for subsequent phylogenetic analysis (described below). Additionally, we identified pseudogenes and partial gene fragments associated with these precursors, leveraging them to deduce the evolutionary process of the new gene. The pairwise alignment of each lineage was visualized using SnapGene software. The alignment was annotated for UTRs, introns, coding sequences, and the amino acid translation (supplementary fig. 5). All main text figures were created with Biorender.

### Phylogenetic Analysis of AFPI and Precursors

In this study, we performed multiple phylogenetic analyses for different purposes. First, to determine whether the *AFPI* evolved independently within each of the three different lineages, we examined the phylogenetic relationships among all *AFPI* family members in the three focal species, winter founder, cunner, and grubby sculpin (supplementary fig. 3). Secondly, to deduce the duplication pattern of gene members in the precursor gene family that facilitated the evolutionary process of the new gene, we reconstructed the phylogenetic trees encompassing all precursor gene family members in *AFPI*+ and *AFPI*- outgroup species within the flounder and wrasse lineage (supplementary fig. 7 and 8). Finally, to infer the evolutionary relationship of the precursor gene, potential intermediate genes, and new genes in the sculpin lineage, we reconstructed their phylogenetic relationships using the sequences from homologous regions (supplementary fig. 9). For each data set, we performed multiple sequence alignment of the nucleotide or amino acid sequences through MUSCLE (Edgar 2004), and phylogenetic analyses using raxmlGUI 2.0 (Edler, et al. 2021). Maximum likelihood phylogenetic trees were constructed using RAxML-NG (Kozlov, et al. 2019) with GTR+FO model of evolution for the nucleotide sequences and the GTR+I+G4 model for the amino acid sequences with 1000 bootstrap replicates. The trees were visualized through the Interactive Tree of Life (ITOL) web-based software (Letunic and Bork 2021). Additionally, a time-calibrated tree (supplementary fig. 1) depicting the divergent times among the species used in this study was constructed using TimeTree5 (Kumar, et al. 2022).

### Protein Structure Prediction of AFPI and Precursors

To explore potential preadaptation for alpha-helical structures within the precursor genes leading to the emergence of the novel AFPI genes, we conducted protein structure predictions. Utilizing AlphaFold2 in a Google Colab notebook (Mirdita, et al. 2022), we predicted the protein structures for the precursor and AFPI genes, as well as any intermediate genes if applicable, for each lineage. The AlphaFold2 model was generated with all default settings, except for the multiple sequence alignment mode, which was changed to a single sequence. The predicted structures were then visualized and annotated using Mol* Viewer (Sehnal, et al. 2021) allowing us to designate the residues that share nucleotide sequence similarity between AFPI and its precursor gene.

## Acknowledgments

We thank Kevin Bilyk for his assistance in acquiring the winter flounder and grubby sculpin specimens. We gratefully acknowledge the support of the Arkansas High Performance Computing Center for providing resources essential to this work. This research was supported in part by the National Institute of General Medical Sciences of the National Institutes of Health under awards P20GM139768, R15GM152956, the Arkansas Biosciences Institute –– the primary research component of the Arkansas Tobacco Settlement Proceeds Act of 2000, and the University of Illinois Foundation Research Fund 339045.

