## supplementary fig. for "Diverse origins of near-identical antifreeze proteins in unrelated fish lineages provide insights into evolutionary mechanisms of new gene birth and protein sequence convergence"

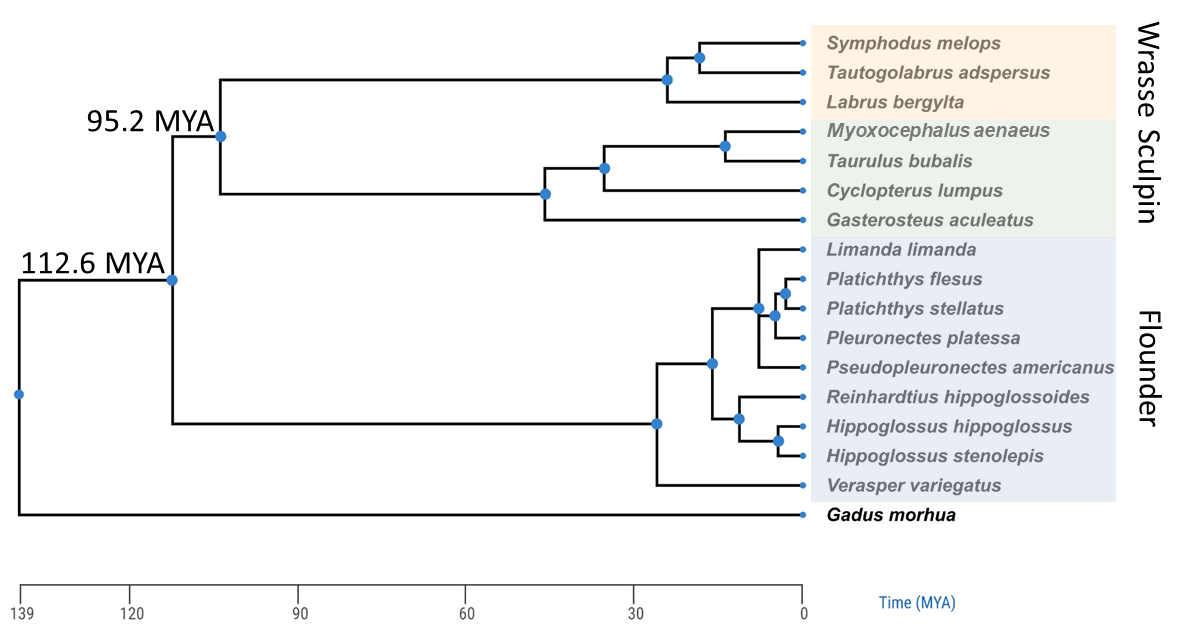

**Supplementary Fig. 1**

**A time-calibrated tree of the species used in this study.** The tree was constructed using TimeTree5 (Kumar, et al. 2022). *Gadus morhua* was included as an outgroup, and *Notolabrus celidotus* (spotty wrasse) was not included as it is not available in the Timetree database. Divergent times between the three taxa (flounder, sculpin, and wrasse) are annotated on the notes.

A) Nucleotide Sequences of AFPI with amino acid translation

**Winter flounder (Pa) AFPI 13**
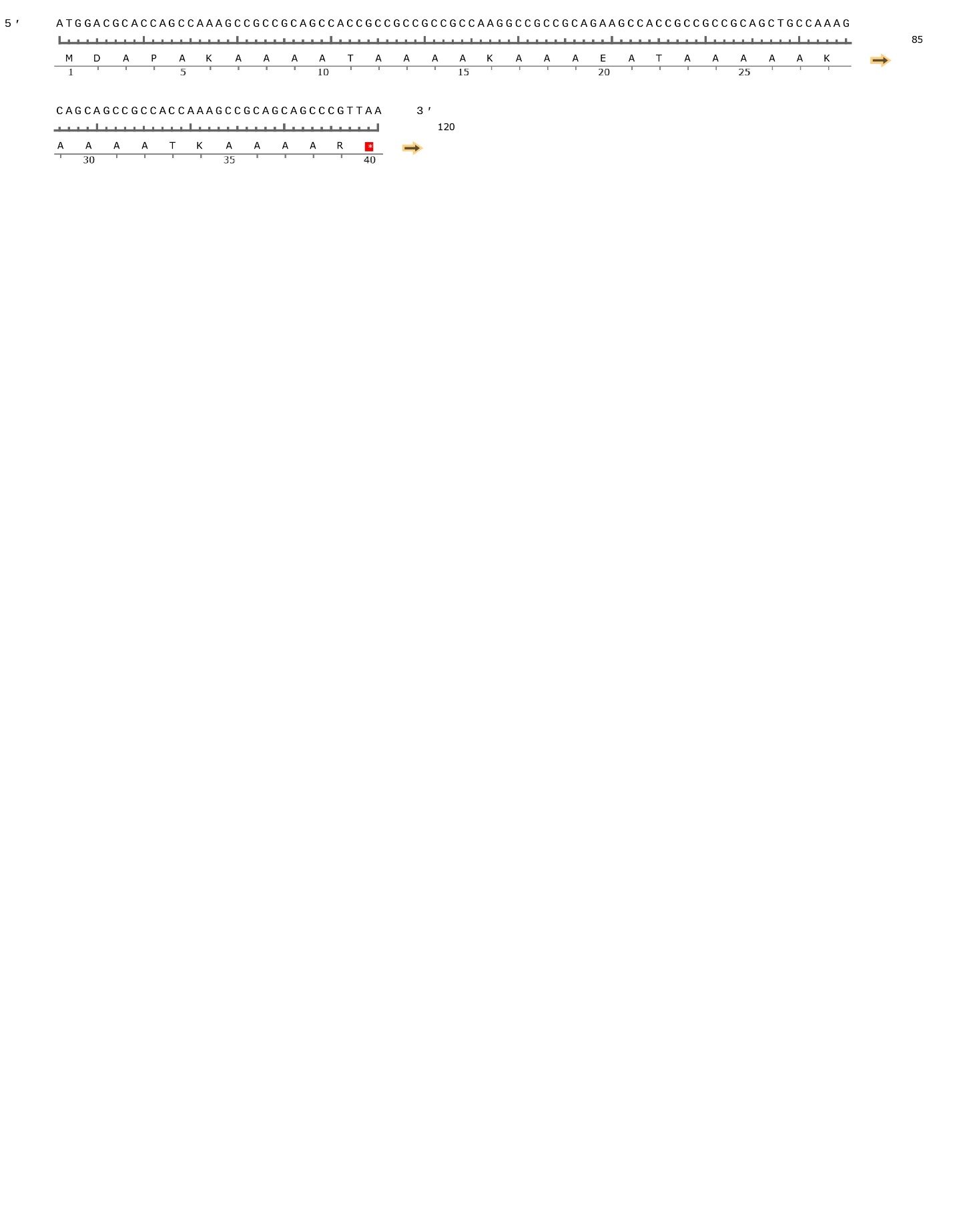

**Cunner (Ta) AFPI 2**
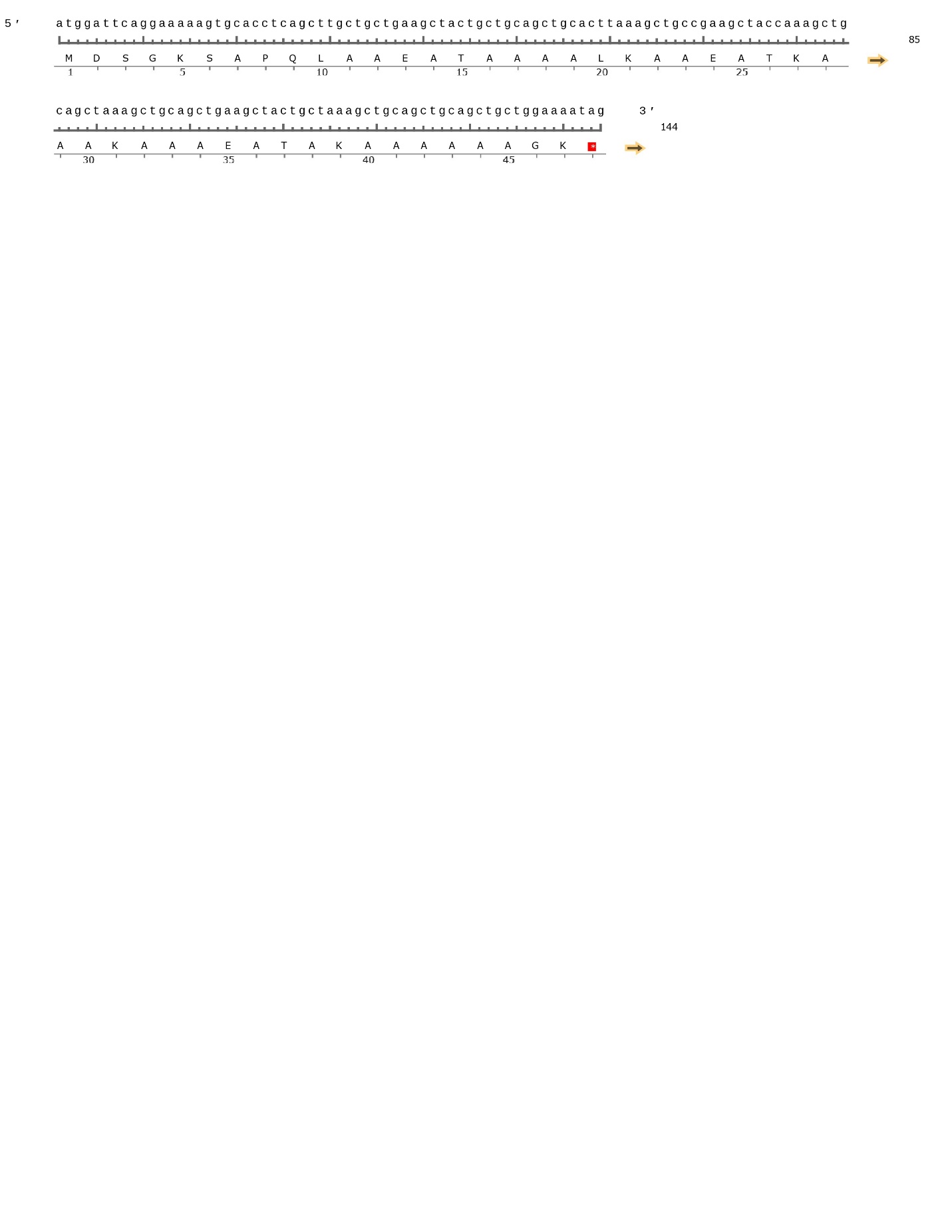

**Grubby sculpin (Ma) AFPI 5**

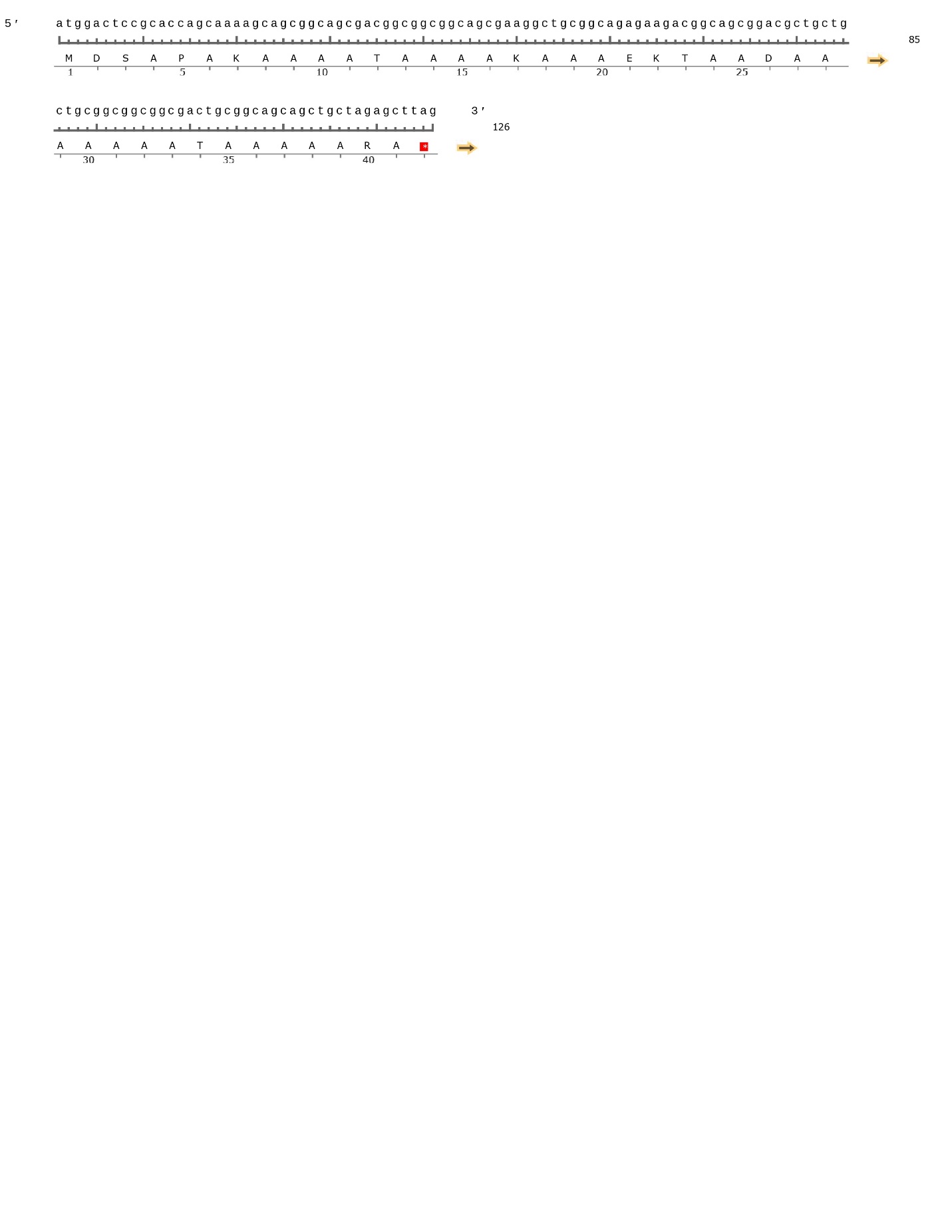

**B)** Amino acid alignment of *AFPI* in all three focal species

```
Ta AFPI 2       MDSGKSAPQLAAEATAAAALKAAEATKAAAKAAAEATAKAAAAAAGK
Pa AFPI 13      MD----APAKAAAATAAAAKAAAEATAAAAAKAAAATKAAAAR----
Ma AFPI 5       MDS---APAKAAAATAAAAKAAAEKTAADAAAAAAATAAAAARA---
                **    **  ** ******  *** * * *  ** **  ***
```

**C)** Nucleotide alignment of *AFPI* in all three focal species

```
Ma AFPI 5       GAAGTTG---------TTGATCTTTCTCTTTTCCAAACGCACCGAGCTAAACAGTAAGTT
Pa AFPI 13      -AGTTTTCATCAGGACTCAAACACTTTTCACTGTCGACCACTCAGGTACGTGAACA----
Ta AFPI 2       CAGATTT------------------------TGTGGA----TCAAGTTCAATAG------
                 *  **                         *    *     *  *      *

Ma AFPI 5       GTCCTTCTATCCATCTGGCTCACACAAAATTCATCTGTTTC-------------AAGGTT
Pa AFPI 13      ----CTCACTTTGTTT--CTTCTATAAATCTGGTTTTACTGTAAATATCTTGGGAAGGAA
Ta AFPI 2       ----TTCTCTCT------CTACAACAAAATGGATTC------------------AGGAAA
                     **  *        **   * ***     *                    * *

Ma AFPI 5       TTGTTGATTTCATGACT-TACCGAAGGACACTTTGACTGAACATTTTTCAAGAATTTTTA
Pa AFPI 13      GGAAGGATATCTGCATTATCCCTGAGGGACCATT---TG-----TTTTACAG--------
Ta AFPI 2       AAGTGAGTGTCTTTATT-TATATGAAGGTATATG---TGATCACTTTTCATG--------
                       * **   * * *     * *     *    **     ****   *


Ma AFPI 5       TAACCTTGCTTTCCTGTTTCAAATTCACCACACTGTTATTATTGTCATACTTAACTCGCA
Pa AFPI 13      ---------CCAGCGGTGAAAGATGAAGATC----------------TTC---------A
Ta AFPI 2       ---------CCTCTTATATGAGATGGATCAC----------------TGC------TGTT
                                *   * **  *   *                * *

Ma AFPI 5       TACATTCTTAACAGTGGTAAA--CTATTCTGCTATCTCATACATGGGATAGTAACATATG
Pa AFPI 13      TCCATGTTCGTCTGATGGAAAGTTTGTTCTGAAACCT-------------TCAGTGGAAG
Ta AFPI 2       TCTTCTCTCGCCCATTGTAGGTGCACCTCAGCTTGCT----------------GCTGAAG
                *      *   *    * *        ** *    **                    * *

Ma AFPI 5       AAACTGCTTTGATTGTATAGCCTTTTATAGCCCATTGAATGCAAACTTGGATTTATATTA
Pa AFPI 13      AAACAGATTCA------------------------TGTCTTCAGGCTTAAACCT------
Ta AFPI 2       CTACTGCTGCAGCTGC----------------------------ACTTAAAGCT------
                  ** * *                                     ***  *  *

Ma AFPI 5       TAGTAAAAAAACAATCTCTCCAAGTAT-----ACAACACTTCAAAATATGTTTGACATTG
Pa AFPI 13      --GCAAAAATCTGAGCTCTGTTAAATCATGGGAAACAACTTTATAATTCAGTCAGGGCTG
Ta AFPI 2       --GCCGAAG--------CTACCAAAGC-------------------TGCAGCTAAAGCTG
                  *   **         **   *                       *           **

Ma AFPI 5       TATATTTCTCTGGAATGAATAGAAAGTGAGAATGGACTCCGCACCAG-CAAAAGCAGCGG
Pa AFPI 13      GAAAACTATTTTATATGCACAGAAGAAGAAGAAGATGTGATCTTTAGTTCATCACCATGG
Ta AFPI 2       CA----------------GCTGAAG-----------------------CTACTGCTAAAG
                 *                   ***                          *   *    *

Ma AFPI 5       CAGCGACGGCGGCGGCAGCGAA--GGCTGCGGCAGAGAAGACGGCAGCGGACGCTGCTGC
Pa AFPI 13      AAACATCATCAGCAGTTAAAGTCTGTCTGCTTCAG------TATCACCGGCCAGTTCCAG
Ta AFPI 2       CTGCAGCTGCAGCTGCTGGAAAATAGCTGCTGGAG------CTGCAGGGGAAGCCGCTGC
                   *  *  * ** *           ****   **         **  **      *

Ma AFPI 5       TGCGGCGGCGGCGACTGCGGCAGCAGCT---GCTAGAGCTTAGTAAAGCAGTT-------
Pa AFPI 13      TGC-TCATG-----TTTCTG-ATCAGCTTGGTTTGAATGATATAAAAACGGATCAAGTCC
Ta AFPI 2       TTCGTC--------CTTTGG-GCCAACT----------------AAAATGGCT-------
                * *  *         *   *   ** **                ***   * *

Ma AFPI 5       ---------------------------GCC------------------------------
Pa AFPI 13      ATGTTTGACCCTGTTTAACACAAGATGGCCACATGGACCATCTTTATTTACATAATGTTT
Ta AFPI 2       -----------------------GCTGGCC-CAGGGA-----------------------
                                           ***

Ma AFPI 5       ---------------TGCTTATAATGCTGGGGAGTGGGGGGG-------------TATTT
Pa AFPI 13      TACATCAGCACTTCCTGTTTCCA--GCCCTAAACTTAAAGAGGCCTCATGGAAACTTCCT
Ta AFPI 2       ---------------TTTTTCCA--GCT----ATTTGGAGGA----TGTGCACACTGTCT
                               *  **  *  **     * *    *               *   *

Ma AFPI 5       GATTATTCG-----ATTTGTTAATCAAACAAAAAGCATCTTGAGACGC------------
Pa AFPI 13      GATGATCTGGTGACACCTGCTGGTTGAAGGAAACAGAGTTTGAGAGGCAGCAGAAAAAAT
Ta AFPI 2       AA------------------------------------CTTGAAATG-------------
                 *                                     **** * *

Ma AFPI 5       --TCCTGTTGTGAATCAG-----CATCATCAATTTAAATGTGTGGT-----TAAAAACCC
Pa AFPI 13      GATTTTAGTTTGAATGAAGAAGCTGTCATTAGATTTTATGTTTGATCACCACACACAGAT
Ta AFPI 2       GACTTCATTGCCAACGAA----------------TGTATGTCTG-------CAAGCATAC
                        *   **  *                 *  **** **        *   *

Ma AFPI 5       G-----------------CTGTTTAG-----ATCTCATAACCAAGAAATGTT----TTTA
Pa AFPI 13      ATTGAACACTGTCATCACTGGGTTCGGTGAAAGTGACGGACCAGTACATGTTGTGATATA
Ta AFPI 2       A-----------------TGAGTTAG--------------------CATGCTATTTTCTA
                                      ** *                     *** *    * **

Ma AFPI 5       CAGCCCGGGAAAAGTGA-----------------ACCTCCACAAAGATCTTT--------
Pa AFPI 13      TAATATTATCATAATAATTATATTAATACCATTAATCTCTGCAGA-ATCACTGACATCAA
Ta AFPI 2       TGTACTTAAAATCATGA-------------------CTTTGCAC--ATCTTTTAC-----
                          *   * *                   **   **   ***  *

Ma AFPI 5       ------------------------------------------------------------
Pa AFPI 13      CATGGACGCACCAGCCAAAGCCGCCGCAGCCACCGCCGCCGCCGCCAAGGCCGCCGCAGA
Ta AFPI 2       ------------------------------------------------------------


Ma AFPI 5       -----------CTTTAG-TGCTAATGTAGTAGCT--------------------------
Pa AFPI 13      AGCCACCGCCGCCGCAGCTGCCAAAGCAGCAGCCGCCACCAAAGCCGCAGCAGCCCGTTA
Ta AFPI 2       -----------CTACGGGTACAAATAAAGTAACC--------------------------
                           *    * * * **   ** * *

Ma AFPI 5       -------------------------------TAAAGA-----------------------
Pa AFPI 13      ATGATCGTGGTCATCTTGATGTGGGATCATGTGAACATCTGAGCAGCGAGATGTTACCAA
Ta AFPI 2       -------------------------------TGAAAGTTT--------------------
                                               * **

Ma AFPI 5       --------------------------------
Pa AFPI 13      TCTGCTGAATAAACCTGAGAAGCTGTTTGTTG
Ta AFPI 2       ------------------GAAACTGTCAA---
                                  *** ****
```

**Supplementary Fig. 2**

**Nucleotide and Amino Acid Sequences from an AFPI gene in the Three Focal Lineages.** A) The nucleotide and translated amino acid sequence of winter flounder *AFPI* 13, cunner *AFPI* 2, and grubby sculpin *AFPI* 5, which were selected for similar gene lengths to improve the alignment quality. B) The amino acid alignment of the three AFPI shows a similar pattern of alanine and threonine. The genes convergently evolved the ice-binding motif that is visually similar in all species. C) The nucleotide alignment of the three AFPI genes shows a dissimilar structure across the gene with low levels of similarity in the coding sequence. Compared to the *AFPI* and precursor gene alignments in supplementary figure 5, the genes are not significantly similar.

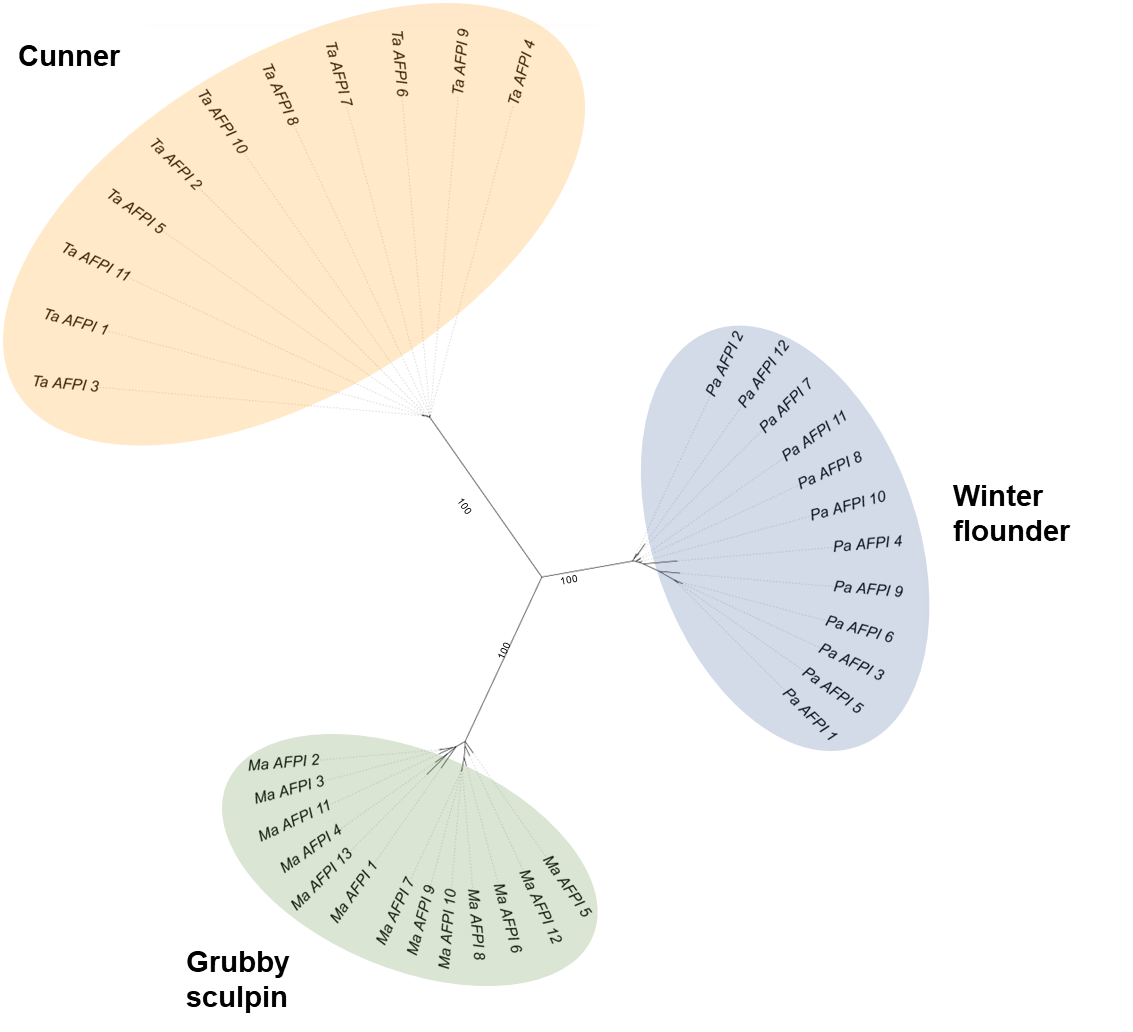

**Supplementary Fig. 3**

**Phylogeny of the AFPI genes in the three focal species winter flounder (*Pseudopleuronectes americanus: Pa*), cunner (*Tautogolabrus adspersus: Ta*), and Grubby sculpin (*Myoxocephalus aenaeus: Ma*).** The numerical designation in the gene names was assigned based on their sequential order within the locus.

**winter flounder**

>Pa AFPI 2

MDAPARAAAA**TAAAAKAAAEATKAAAAKAAAATKAAA**H*

>Pa AFPI 3

MALSLFTVGQLIFLFWTMRITEARPDPAAKAAPAAAAAPAAAAPD**TASDAAAAAALTAANAKAAAELTAANAAAAAAATA**RG*

>Pa AFPI 4

MSAPARAAAATAATAKAAAEA**TAADAAAAAALTAATAAAAAAATAANAAAAAAATIEAAAAAAAATAAGATAAADAAAAAAAATAAAAAAAAKATIDDAAAAAATTAANAAAAAAATAATAAAAAAATKATIDDAAAAAAATAATAAAAAAATAATAAAAAAATIKAAAAAAAATAAAA**C*

>Pa AFPI 5

MALSLFTVGQFIFLFWTIRITEANIDPAARAA**TAAAASKAAVTAADAAAAAATIAASAASVAAATAADDAAASIATINAASAAAKSIAAAAAMAAKDTAAAAASAAAAAVASAAKALETINVKAAYAAATTANTAAAAAAATATTAAAAAAAKATIDNAAAAKAAAVATAVSDAAATAATAAAVAAATLEAAAAKAAATAVSAAAAAAAAAIAFAAAP***

>Pa AFPI 6

MAASLFTVGQIIFLFWTIRITEARPDPAAKAAPAAAAVPAAAAPD**TASDAAAAAALTAANAAAAAKLTADNAAAAAAATA**RG*

>Pa AFPI 7

MALSLFTVGQLIFLFWTMITEARPDPAAKAAPAAAAAPAAAAPD**TASDAAAAAALTAANAKAAAAKLTAANAAAAAAAATA**RG*

>Pa AFPI 9

MAHSLFTVGQFIFLFWTIRITEANINPAARAAAAAAAAQAAA**TAADAAAAAATIAATAATIAAATAADVAAASIATINCASAAAKSIAAATAMAAEDTAAAAASAAADTVAAAAKALETINVKAAYAAATTANTAAAAAAATATTAAAAAATKATIDNAAAAKAAAVATAVAAAAATAATAAAVAAATLGAAAAKAAATAVSAAATAAAAAIAFAAA**P*

>Pa AFPI 10

MDAPAKAAAA**TAAAAKAAAEATAAAAAKAAAATKAAAA**R*

>Pa AFPI 12

MDAPAKAAAA**TAAAAKAAAEATAAAAAKAAAATKAAAA**R*

>Pa AFPI 13

MDAPAKAAAA**TAAAAKAAAEATAAAAAKAAAATKAAAA**R*

>Pa AFPI 14

MDAPAAAAAA**TAAAAKAAAEATAAAAAAAAAATAEAAAKAAAATKAAAAAAAA**R*

**grubby sculpin**

>Ta AFPI 1

MDSRKSAPELAAEA**TAAAAAKAARDTAEAAAKAAEATRAAAAAAAEAAAKAAEATKAAAKAAAEATARAAAAAA**GK*

>Ta AFPI 2

MDSGKSAPQLAAEA**TAAAALKAAEATKAAAKAAAEATAKAAAAAA**GK*

>Ta AFPI 3

MDSRKSAPELAAEA**TAAAAAKAARDTAEAAAKAAEATRAAAAAAAEAAAKAAEATKAAAKAAAEATARAAAAAA**GK*

>Ta AFPI 4

MDSGKSAPQLAAEA**TAAAALKAAEATKAAAKAAAEATAKAAAAAA**GK*

>Ta AFPI 5

MDSGKSAPQLAAEA**TAAAALKAAEATKAAAKAAAEATAKAAAAAA**GK*

>Ta AFPI 6

MDSGKSAPQLAAEA**TAAAALKAAEATKAAAKAAAEATAKAAAAAA**GK*

>Ta AFPI 7

MDSGKSAPQLAAEA**TAAAALKAAEATKAAAKAAAEATAKAAAAAA**GK*

>Ta AFPI 8

MDSGKSAPQLAAEA**TAAAALKAAEATKAAAKAAAEATAKAAAAAA**GK*

>Ta AFPI 9

MDSRKSAPQLAAEA**TAAAALKAAEATKAAAKAAAEATAKAAAAAA**GK*

>Ta AFPI 10

MDSGKSAPQLAAEA**TAAAALKAAEATKAAAKAAAEATAKAAAAAA**GK*

>Ta AFPI 11

MDSRKSAPELAAEA**TAAAAAKAARDTAEAAAKAAEATRAAAKAAAEATARAAAAAA**GK*

**cunner**

>Ma AFPI 1

MDAPARAAAAAAAAAAAAA**TAAAATAAAAAEAAATTAANAAAAAAATAAAAATAATAAEATAAEAAEAAAATAAEAAEAAATTAANAAAAAAATAAAAAAAAEATAAAAAAAAAEAAEAAAATAAAAAAAA**V*

>Ma AFPI 2

MDAPARAAAAAAA**TAAEAAALTAATAAAAAAATAANAAAAAAATANTAAAAAALAAANAKAAAAA**V*

>Ma AFPI 3

MDAPARAAAAAAA**TAAEAAALTAATAAAAAAATAANAAAAAAAAATANTAAAAAALAAANAKAAAAA**V*

>Ma AFPI 4

MDAAAK**TKAAAAAAAAATVAAAAAAAAETAAAANKAAEAAAEAAAKVAEAAAKAAAATAAAAAAAAA**V*

>Ma AFPI 5

MDSAPAKAAAA**TAAAAKAAAEKTAADAAAAAAATAAAAARA***

>Ma AFPI 6

MDAPAIAAAK**TAADALAAANKTAADAAAAAAKTA**GK*

>Ma AFPI 7

MDAPAIAAAK**TAADALAAAKKTAADAAAAAAKTA**GK*

>Ma AFPI 8

MDAPAIAAAK**TAADALAAAKKTAADAAAAAAKTA**GK*

>Ma AFPI 9

MDAPAIAAAK**TAADALAAAKKTAADAAAAAAKTA**GK*

>Ma AFPI 10

MDAPAIAAAK**TAADALAAAKKTAADAAAAAAKTA**GK*

>Ma AFPI 11

MEAAAAAKAAA**TAAANAAAKAAAAAIAEAAEAAEAAEAAATKAANAAAEAAAKSAAAAAEAKANAAAAAAAAAAAAAAAAA***

>Ma AFPI 13

MDAPARTAASR**TKAAKEAAAARAAAMKAAAA**N*

**Supplementary Fig. 4**

**The amino acid sequences of all intact AFPI in the three focal species.** The repetitive alanine-rich region is shown in blue, with all the threonine residues underlined. The shared N-terminal motif between winter flounder (Pa) and grubby sculpin (Ma) is highlighted in yellow. The shared C-terminal residues between cunner (Ta) and grubby sculpin (Ma) are highlighted in pink. Gene naming follows supplementary figure 3, and pseudogenes were not included in this figure.

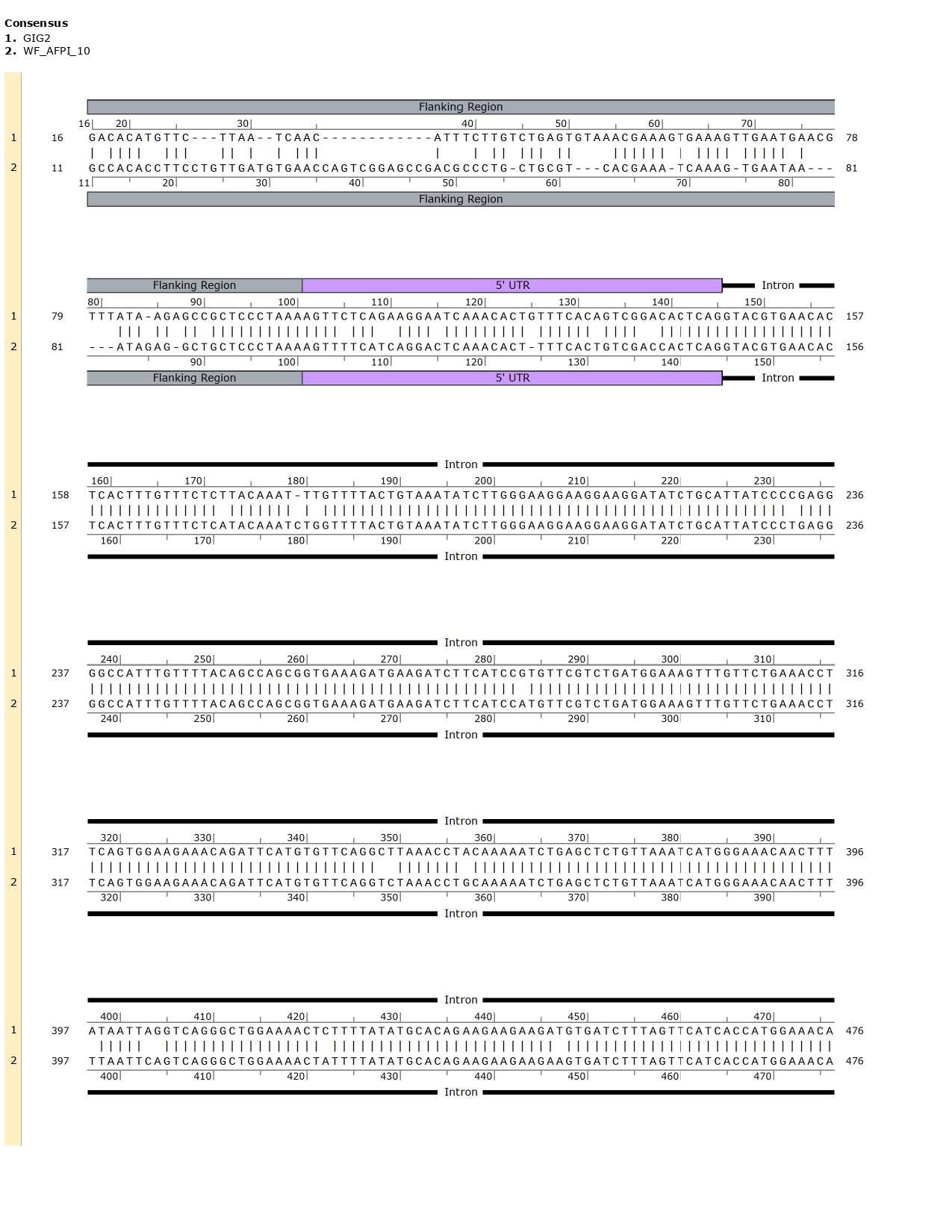

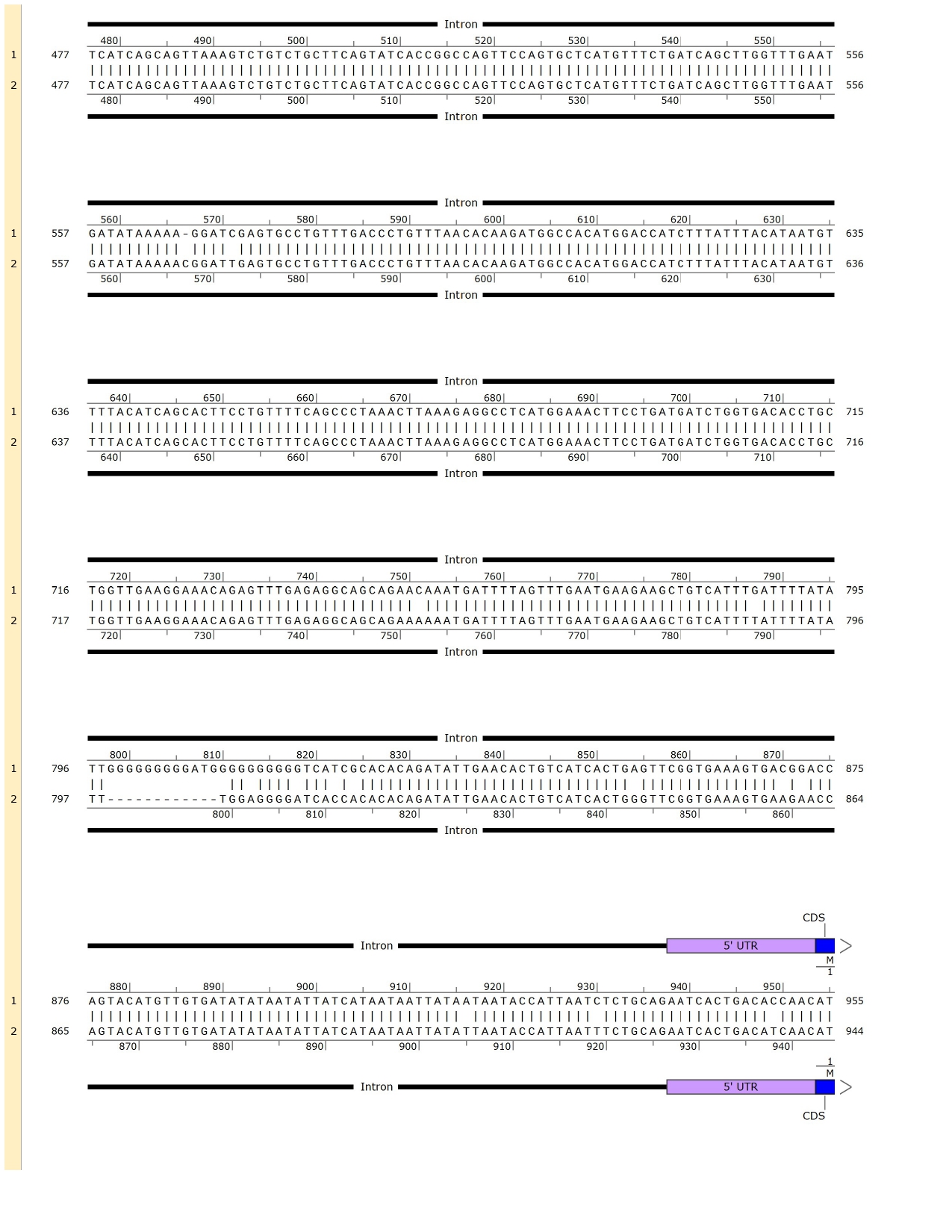

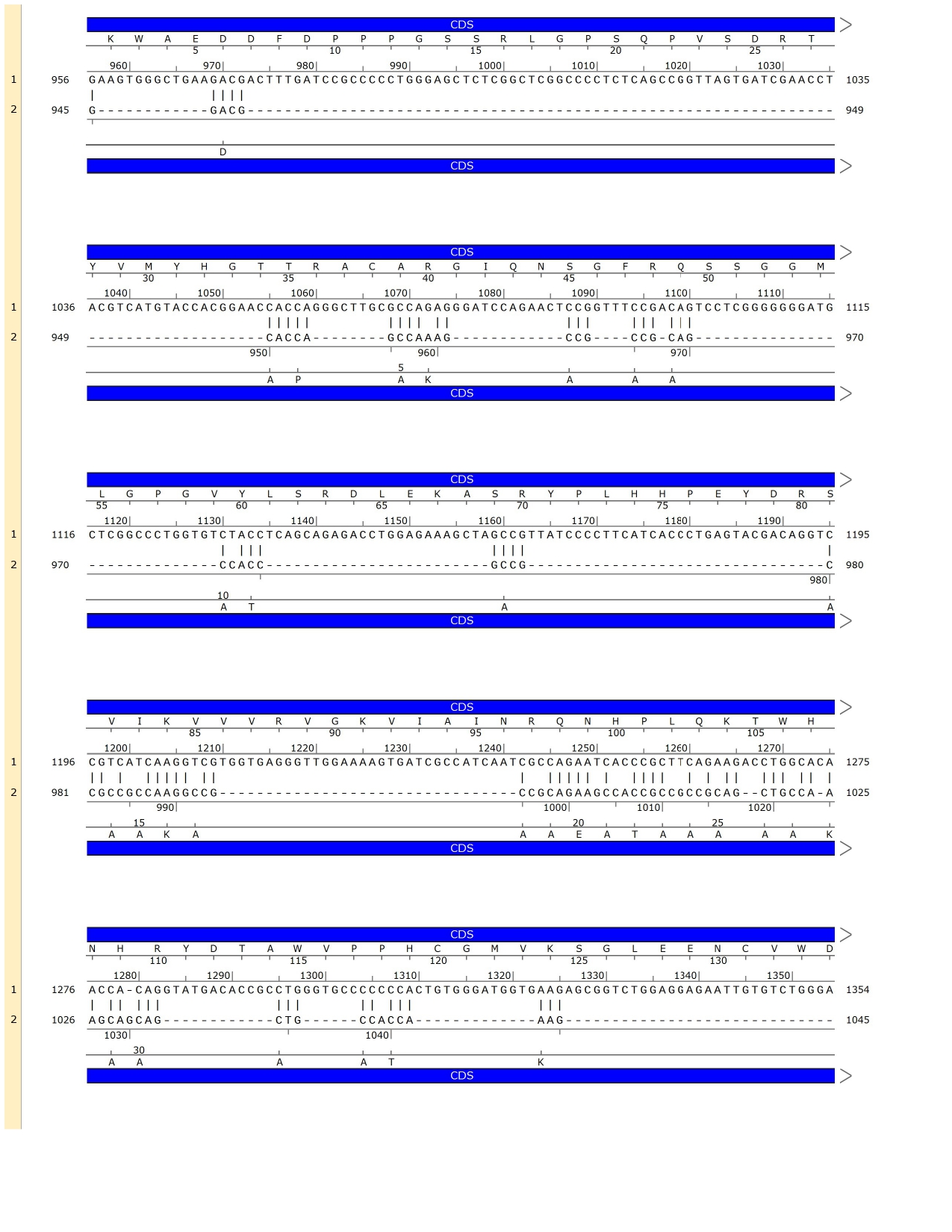

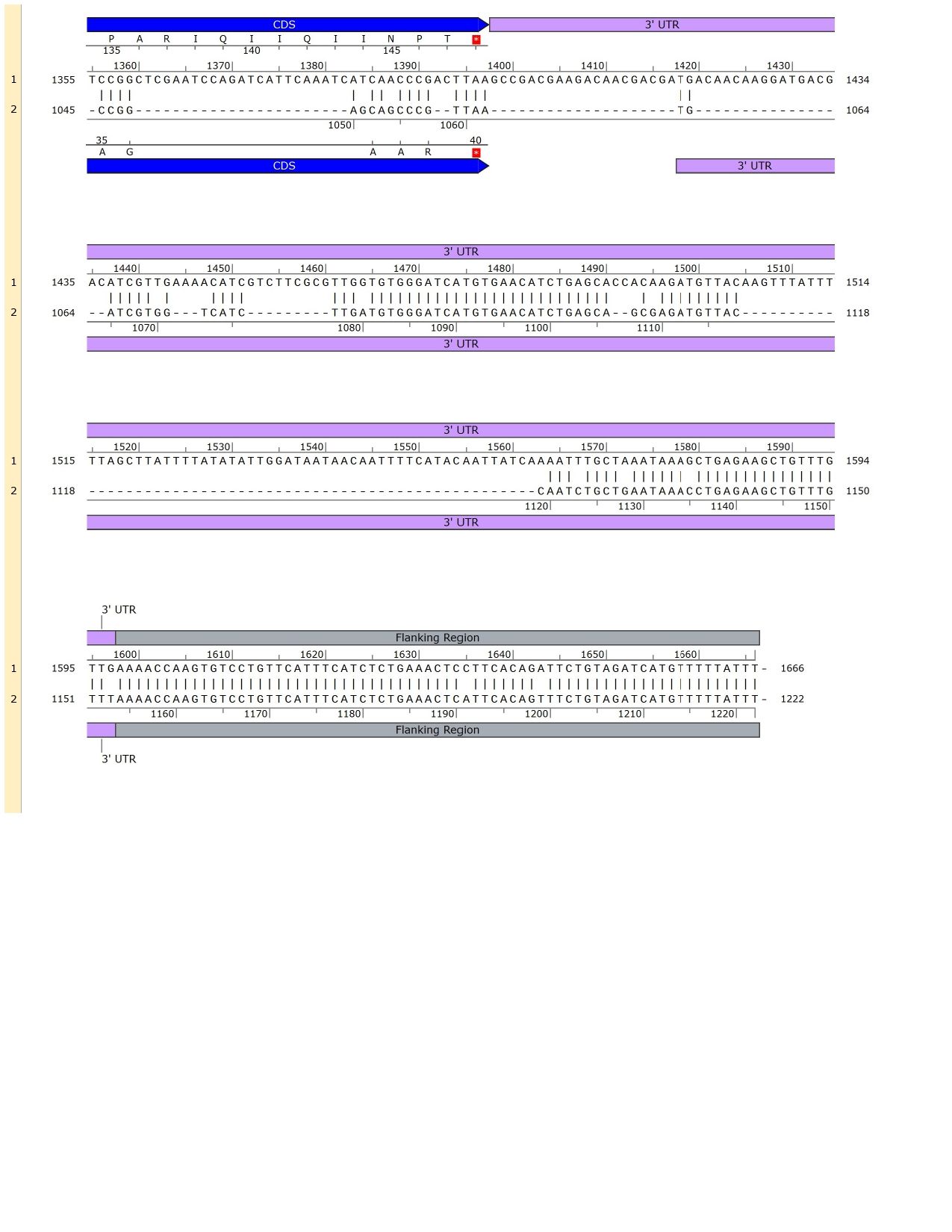

***GIG2* Aligned with *AFPI* 10 in Winter Flounder**

1. *GIG2*
2. *AFPI*

A)

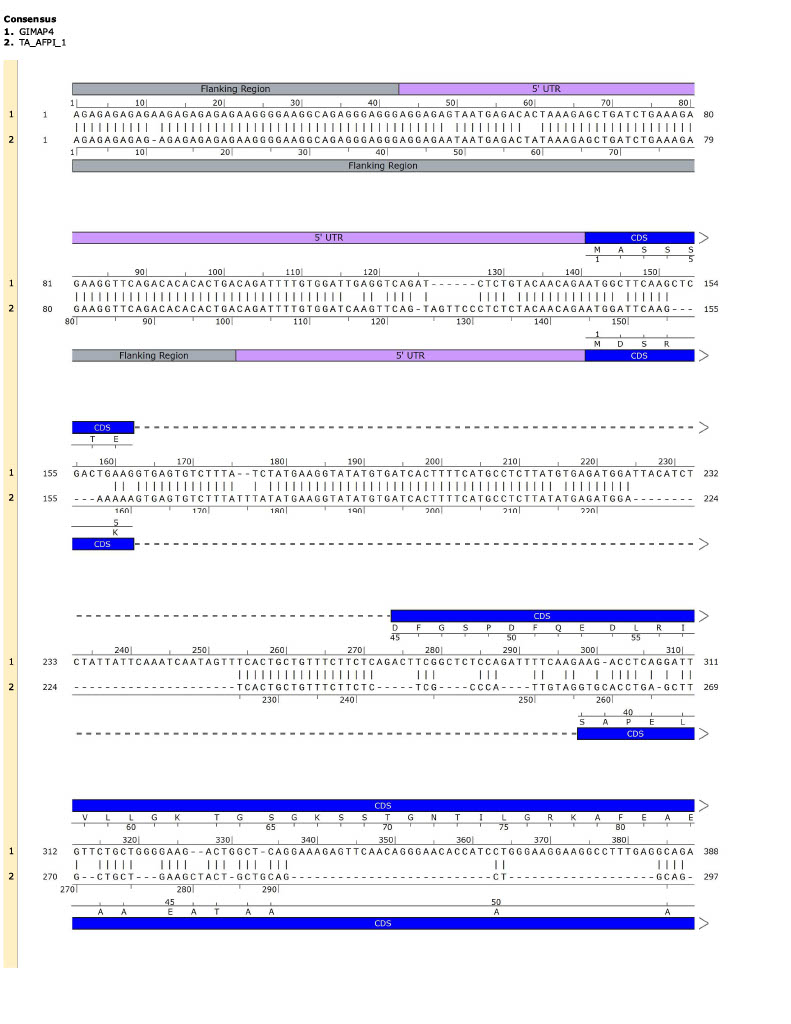

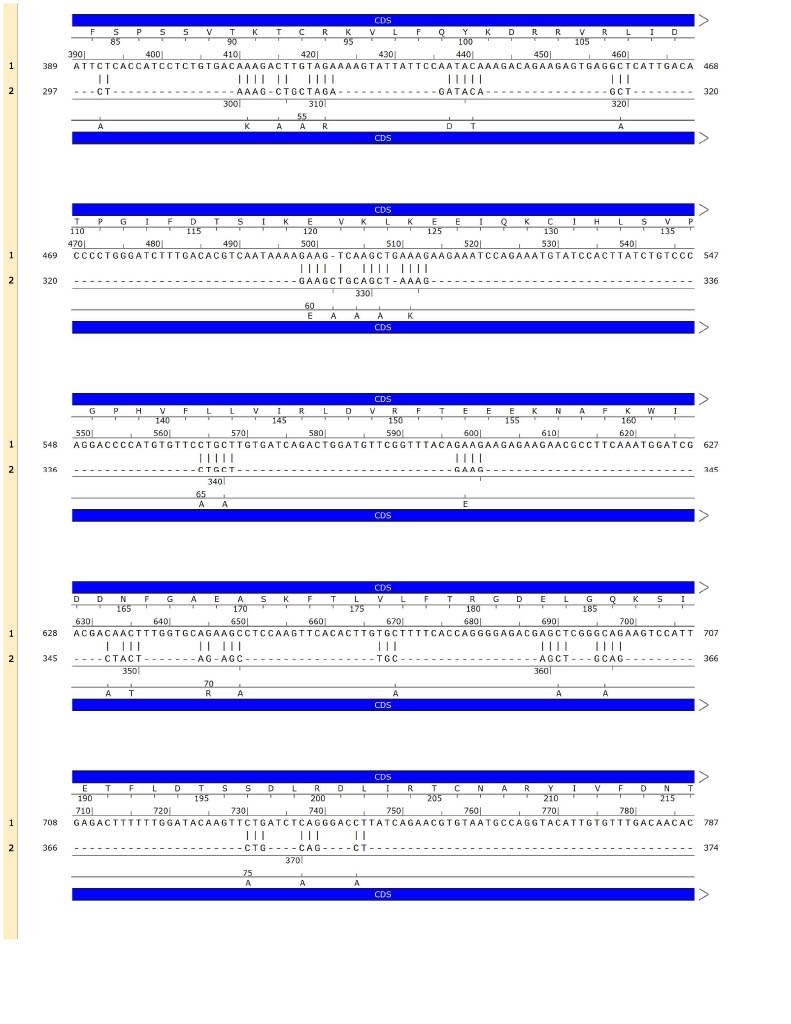

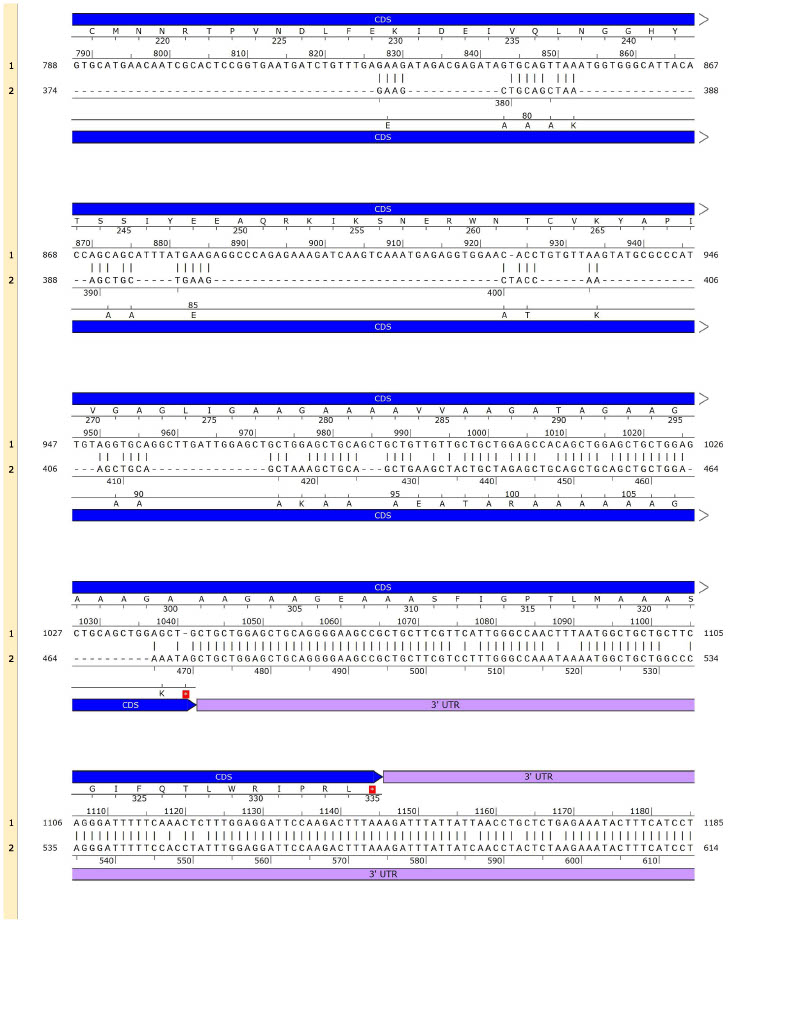

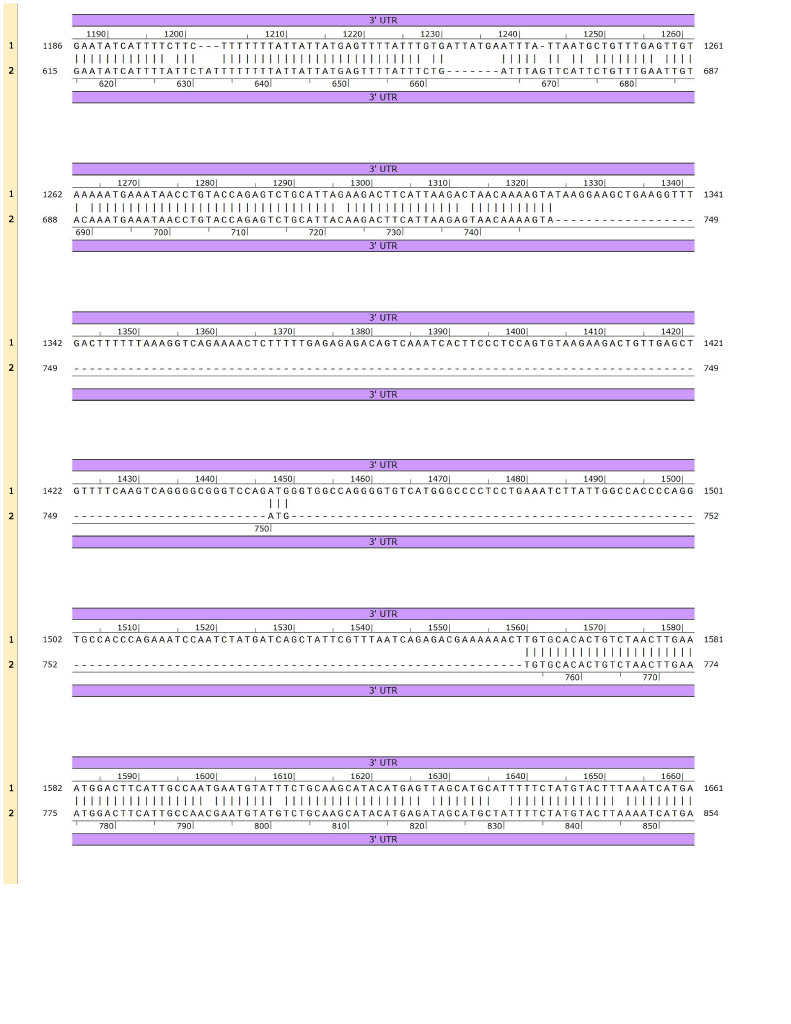

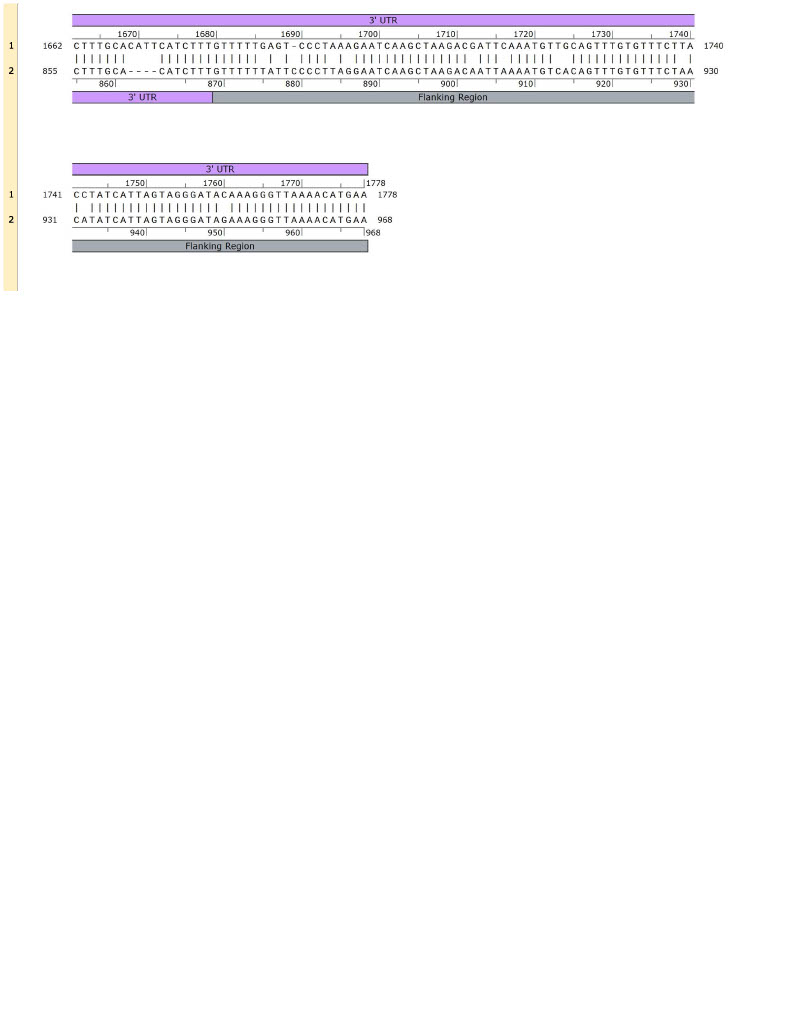

B)

***GIMAP4* Aligned with *AFPI* 1 in Cunner**

1. *GIMAP4*

2. *AFPI*

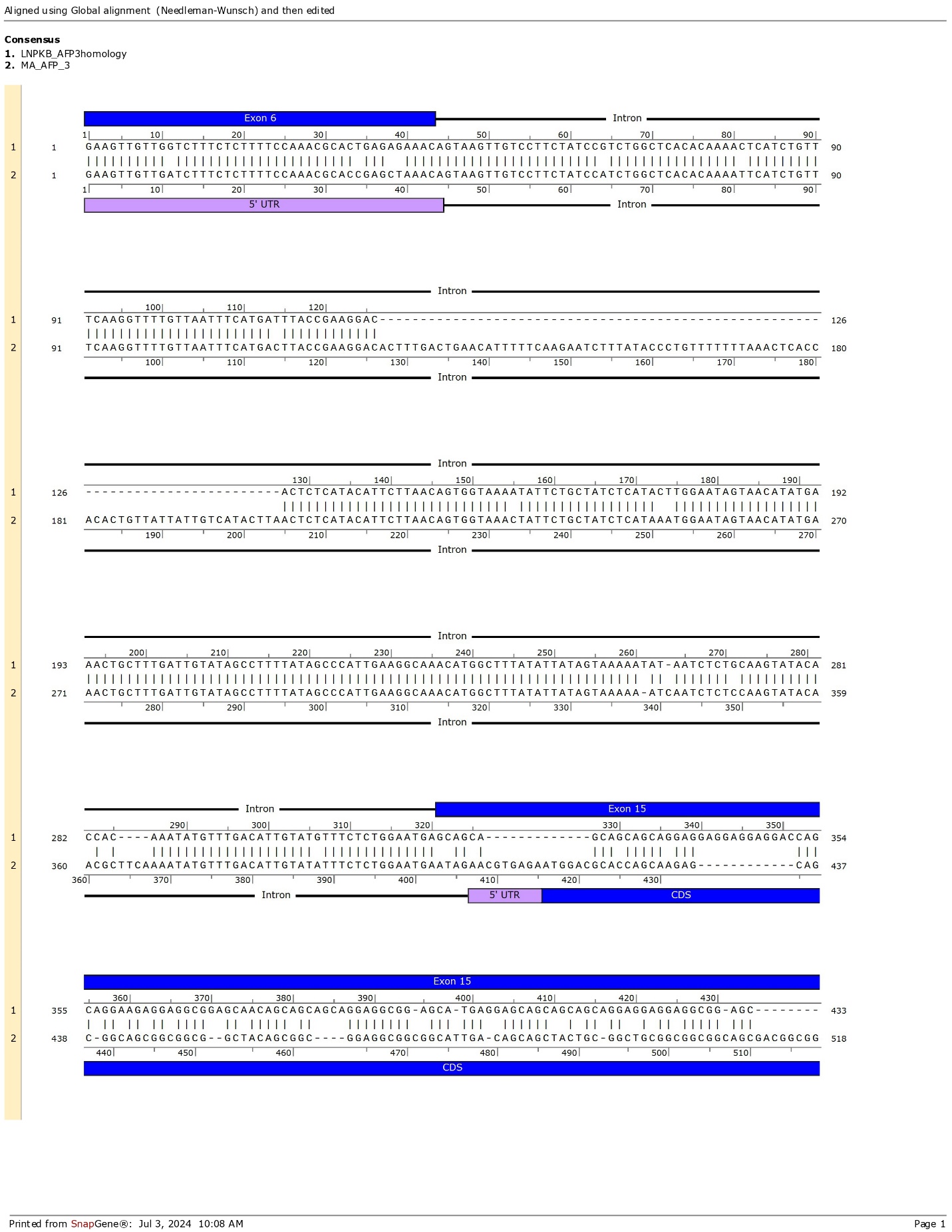

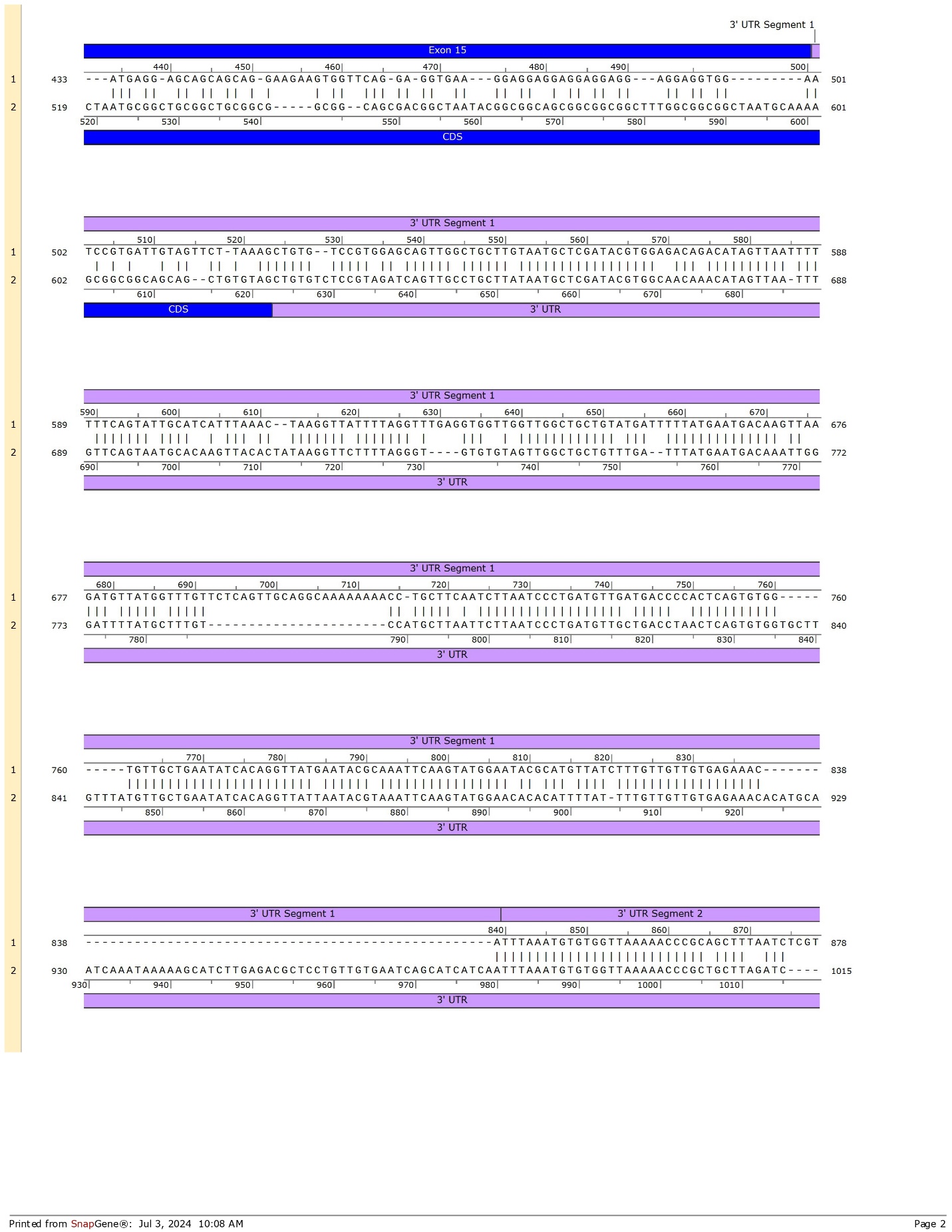

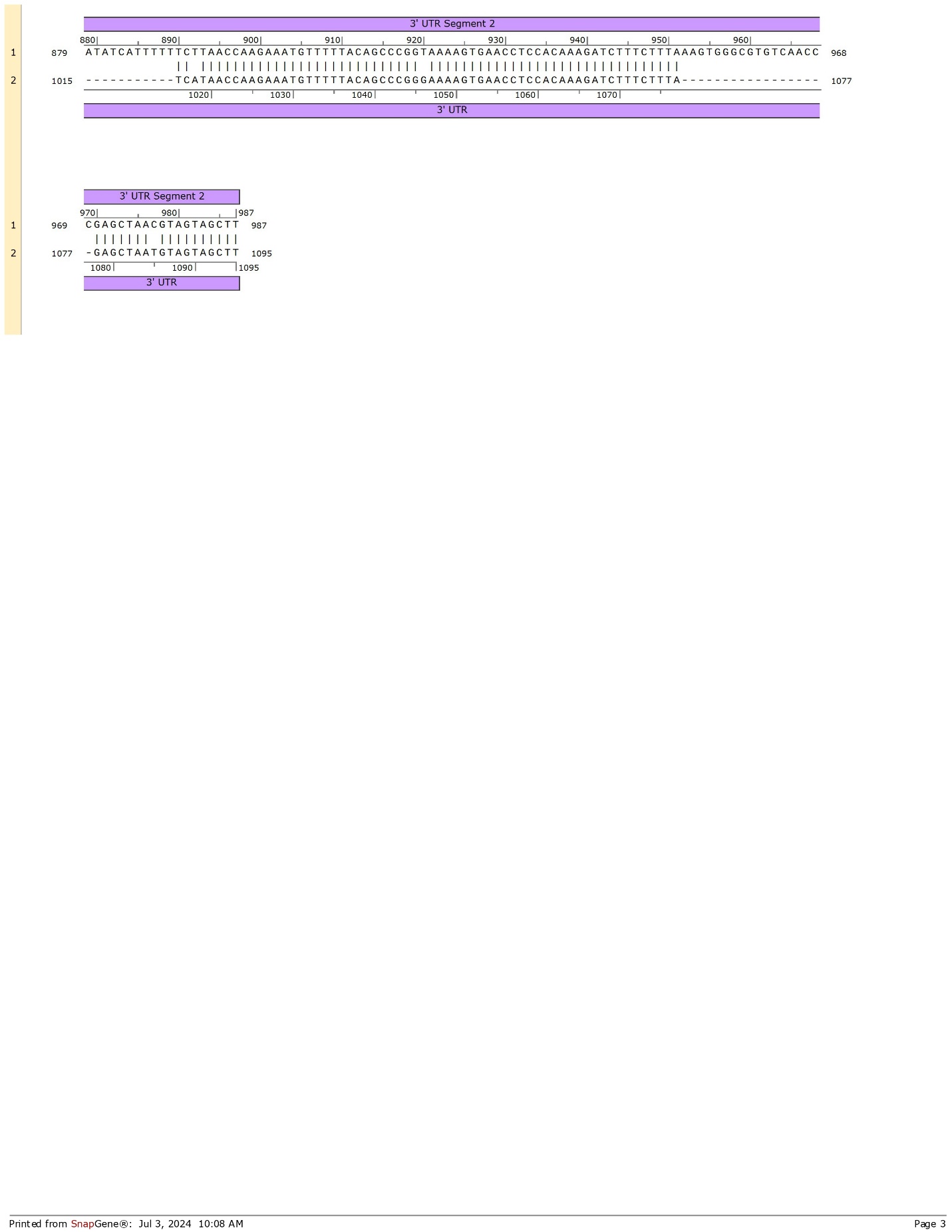

4251 NT

A

G

C)

***LNPKB* Aligned with *AFPI* 3 in Sculpin**

1. *LNPKB* Homologous Regions

2. *AFPI*

384 NT

A

385 NT

A

T

A

**Supplementary Fig. 5**

**DNA sequence alignment between *AFPI* and its respective precursor gene in each focal species**. The pairwise alignments were performed and visualized in SnapGene to show the nucleotide alignment, amino acid sequence, and annotated features for UTRs, exons, introns, and coding sequences. Annotations for AFPI genes were based on the following annotated genes: GenBank: X53718.1 (winter flounder), GenBank: JF937681.2 (cunner), and GenBank: MH745497.1 (sculpin). The ancestral gene annotations were derived from GenBank: UUW46980.1 (*GIG2*), XM_020654175.2 (*GIMAP4*), and XM_034551604.1 (*LNPKB*). For *LNPKB*, only the regions sharing sequence identity with AFPI are shown. Regions without sequence identity are omitted, with their lengths indicated in nucleotides (NT).
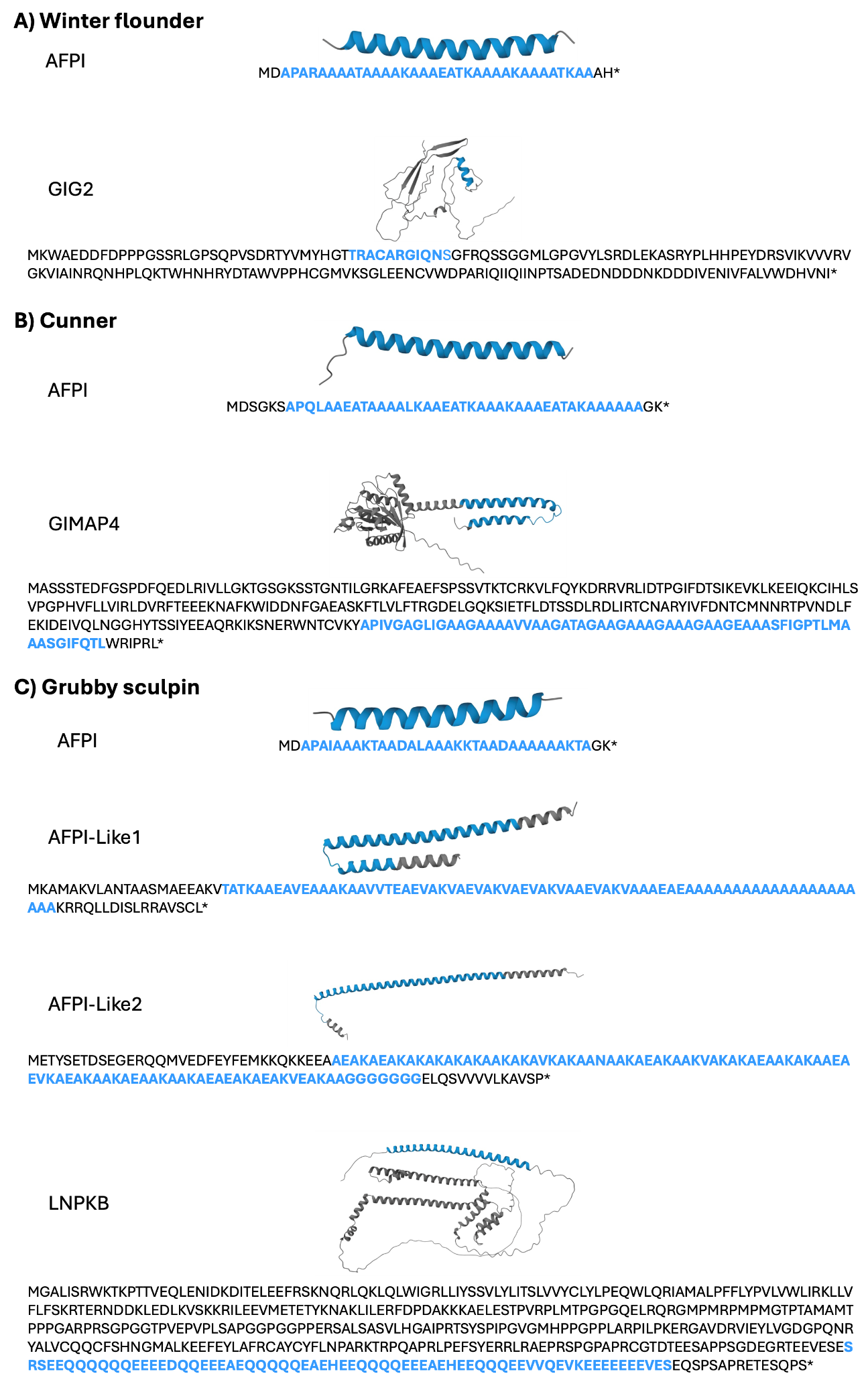

**Supplementary Fig. 6**

**Protein structure predictions for AFPI and its precursor in the three focal species.** For AFPI, the amino acid sequences containing an alanine-rich repetitive region with evenly spaced threonine and the analogous area of the predicted protein structure are depicted in blue. For the precursor and AFPI-like proteins, the highly repetitive region with coding sequence similarity is indicated in blue. The AFPI and precursor proteins shown here are winter flounder Pa_AFPI_2, Ta_GIG2_1, cunner Ta_AFPI_2, Ta_GIMAP4_1, and grubby sculpin Ma_AFPI_6.

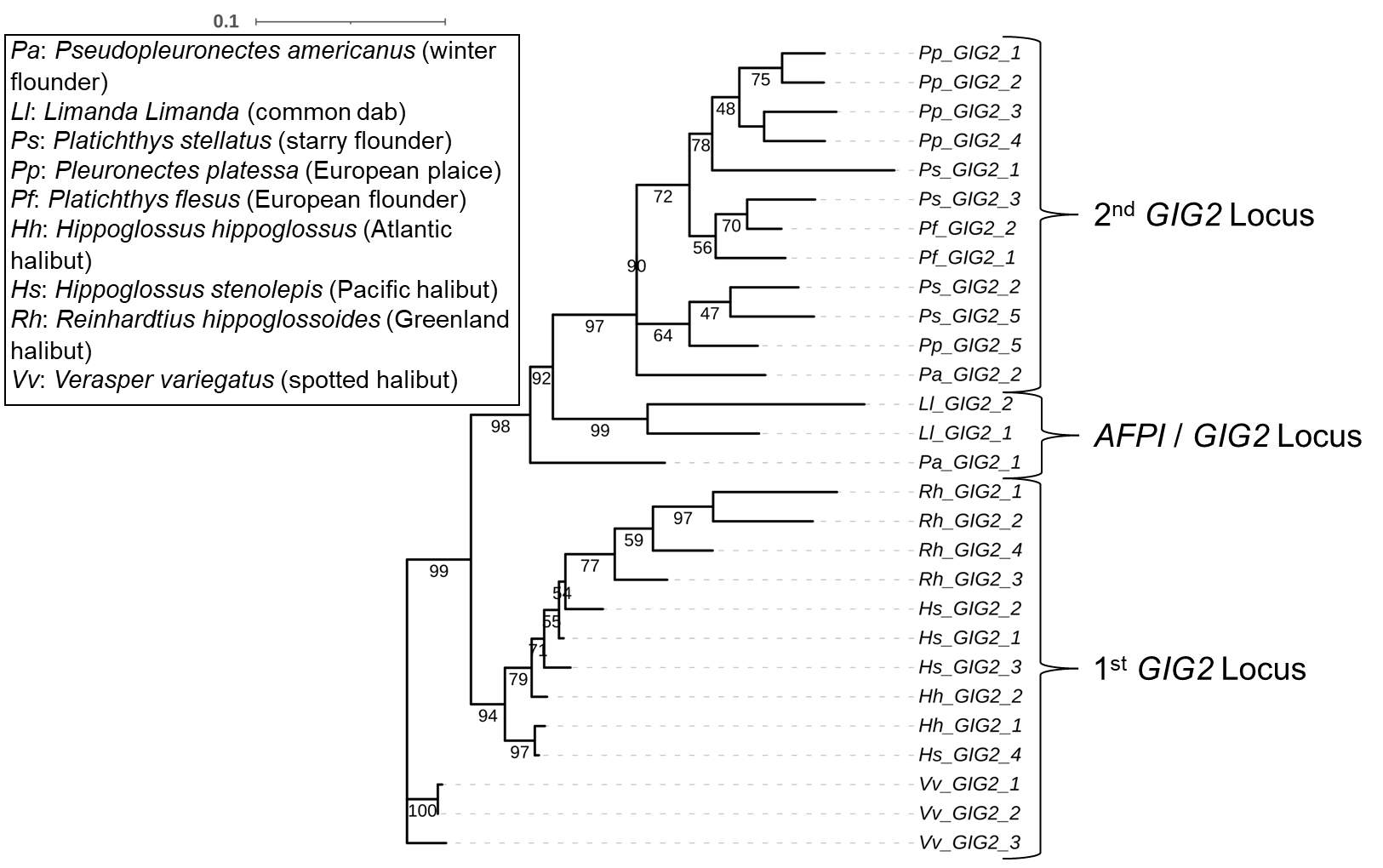

**Supplementary Fig. 7**

**Maximum likelihood tree for all GIG2 genes in AFPI+ and AFPI- species in the flounder lineage.**

The numerical designation in the gene names was assigned based on their sequential order within their respective locus as shown in Fig. 4. Bootstrap values (%) are labeled at branch nodes.

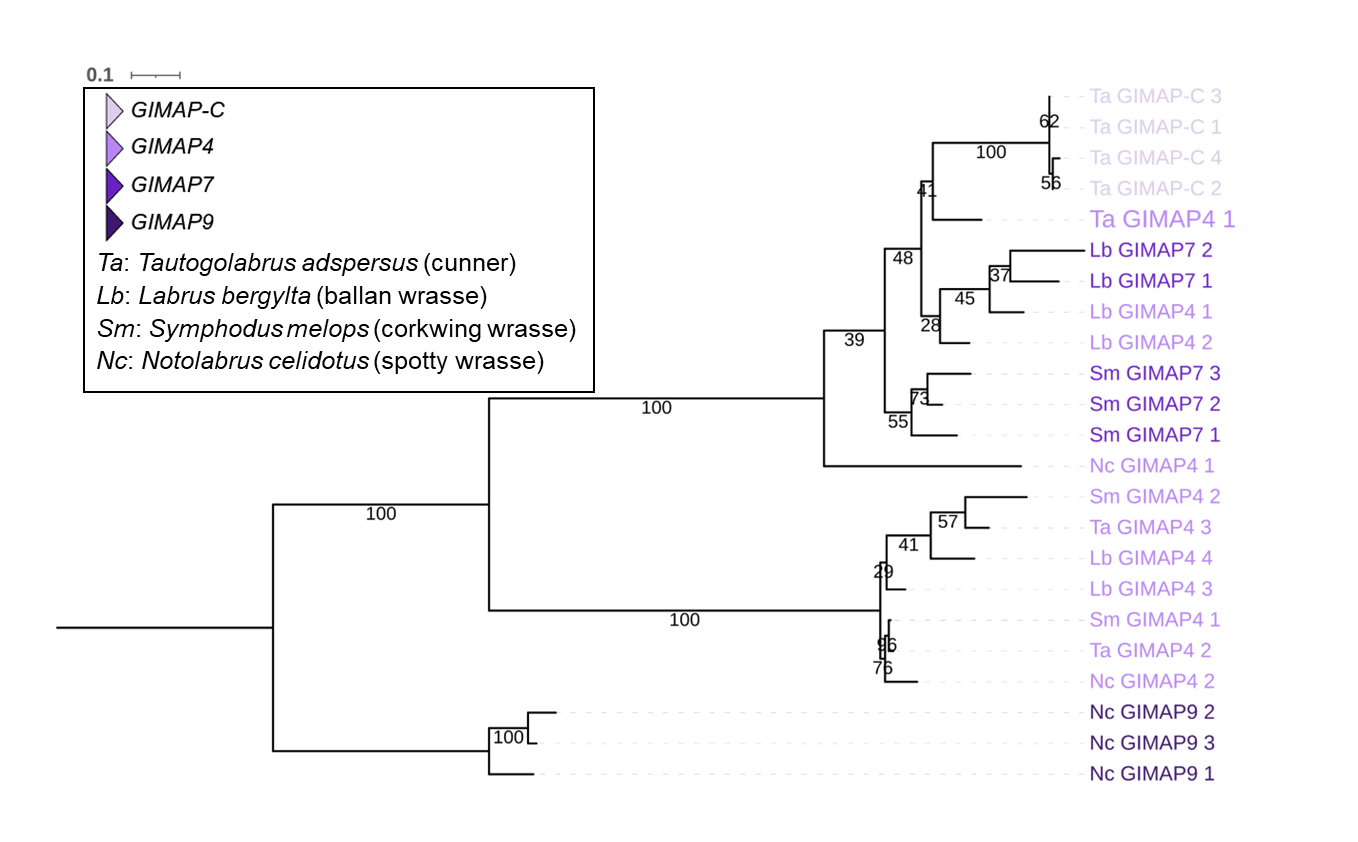

**Supplementary Fig. 8**

**Maximum likelihood tree for GIMAP amino acid sequences in the wrasse lineage.** The text color corresponds to the *GIMAP* locus maps shown in Fig. 5. The numerical designation in the gene names was assigned based on their sequential order within their respective locus as shown in Fig. 5. The precursor gene showing the highest sequence identity with *AFPI* is highlighted in larger font size. Bootstrap values (%) are labeled at branch nodes. The tree was constructed with amino acid sequences rather than nucleotide sequences due to the presence of extensive indels and repeats in the nucleotide sequence, which led to challenges in aligning homologous sequences.

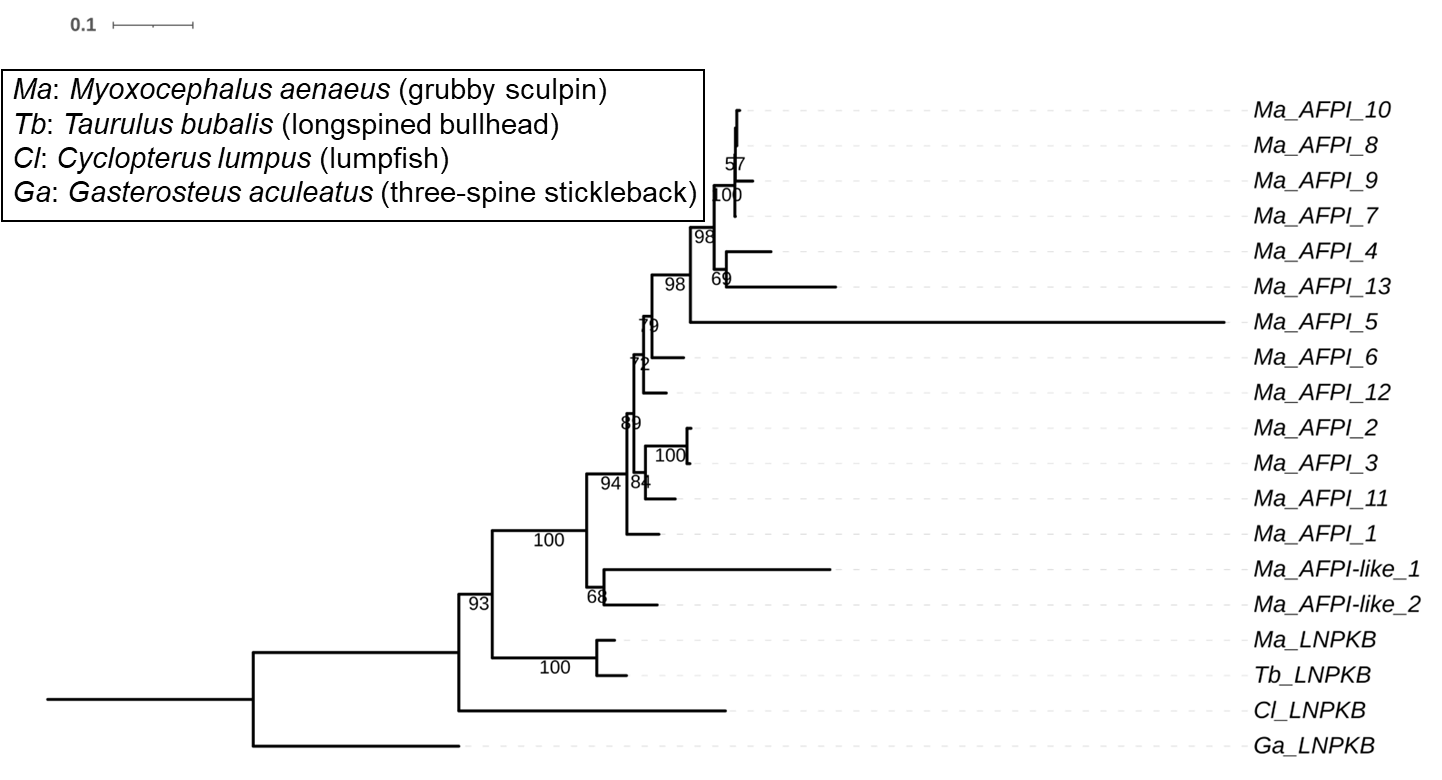

**Supplementary Fig. 9**

**Maximum likelihood tree of *AFPI* and *AFPI-like* genes, the precursor gene *LNPKB* in grubby sculpin (*Ma*), and from related *AFPI-* species (*Tb, Cl, Ga*).** *AFPI* gene names correspond to Supplementary Fig. 3. Bootstrap values (%) are labeled at branch nodes.
